# CSR-1 RNA interference pathway restricts holocentromere protein CENP-A/HCP-3 localization in *Caenorhabditis elegans*

**DOI:** 10.1101/2022.05.26.493264

**Authors:** Charmaine Yan Yu Wong, Karen Wing Yee Yuen

## Abstract

CSR-1 is an argonaute of a RNA interference pathway that is important for chromosome segregation in *C. elegans*. Live-cell imaging revealed that CSR-1 depletion slows down spindle pole separation in a kinetochore-dependent manner. In *csr-1*(RNAi) embryos, the kinetochores may be misattached to the microtubules and chromosome segregation is disrupted. On the holocentromeres, there are increased levels of some kinetochore proteins, including the centromeric epigenetic mark, CENP-A or HCP-3. Without affecting HCP-3 expression level, HCP-3 density is higher on stretched chromatin fibers in CSR-1-depleted embryos. The increased HCP-3 deposition on chromatin after CSR-1 depletion is at least partially independent of HCP-3 loading factors, KNL-2 and LIN-53, suggesting a non-classical, improper HCP-3 loading pathway. Negative regulation of HCP-3 holocentromere loading by CSR-1 required its slicer activity and the b isoform. CSR-1 acts as a HCP-3 repressor for its chromosomal occupancy, shedding light on the role of RNAi pathways in specifying the localization of centromere proteins.

## Introduction

Equal segregation of chromosomes during cell division is crucial for an organism’s survival. Microtubules attach to chromosomes at the centromeres to pull them apart. During mitosis, the sister chromatid centromeres are bi-oriented, each facing toward the opposite spindle poles (Tanaka, 2002). One of the functions of centromere is to assemble kinetochore, so that microtubules can bind. While this function is conserved among organisms, centromere architecture and centromeric DNA sequences vary greatly. In contrast, CENP-A, a histone H3 variant, is found on the active centromeres in many organisms, even on ectopic centromeres that are known as neocentromeres (Amor and Choo, 2002, De Rop et al., 2012). CENP-A is at the top of the hierarchy for centromere identity and kinetochore assembly. Depletion of CENP-A leads to kinetochore-null (knl) phenotypes and severe chromosome missegregation (Oegema et al., 2001).

In many organisms, CENP-A is associated with repetitive sequences at the monocentromere on the chromosome, for example, α-satellite DNA in humans and the minor satellite DNA in mice (Guenatri et al., 2004, Sullivan et al., 2017). Holocentromeres are positioned along the entire poleward faces on the replicated, condensed sister chromatids (Buchwitz et al., 1999). Interestingly, the number of microtubules attached per chromosome is similar for human centromeres and *C. elegans* holocentromeres (McEwen et al., 1998, Redemann et al., 2017). HCP-3, the CENP-A homolog in *C. elegans*, is found to be associated to about 25-50% of the genome, which included both genic and inter-genic DNA regions (Gassmann et al., 2012, Lin et al., 2021). HCP-3-binding locations seem to be influenced by transcription. By cross-linked chromatin immunoprecipitation-DNA microarray chip (chIP-chip), HCP-3 occupancy is inversely correlated to RNA polymerase II occupancy in embryos (r= -0.67) (Gassmann et al., 2012). HCP-3 occupancy is correlated positively with repressive histone modifications H3K27me3 (r= 0.64) and negatively with H3K36me3 (r= -0.6). H3K27me3 is a repressive mark whereas H3K36me3 is associated with active transcription. In another native chIP-seq study, HCP-3 localizes to about 700 nucleosome-sized peaks with low nucleosome turnover. These peaks coincide with high occupancy target (HOT) sites, which may be occupied by transcription factors in somatic cells (Steiner and Henikoff, 2014). Although RNAPol Il is no longer present at these sites in the embryos, regions that are only transcribed in the germline (169 genes) surprisingly also have low HCP-3 occupancy in embryos (average z-score <-0.5) (Gassmann et al., 2012).

Germline transcripts are the major targets for a germline RNA interference (RNAi) pathway containing argonaute CSR-1, short for chromosome segregation and RNAi-deficient-1. CSR-1-bound short interference RNAs (siRNAs) are antisense to over 4000 germline-expressed genes (Claycomb et al., 2009, Gu et al., 2009). These CSR-1-associated small RNAs are primarily 22 nucleotides in length, with 5’ Guanosine triphosphate, and therefore are called the 22G-siRNA. The CSR-1-bound small RNAs are synthesized by the RNA-dependent RNA polymerase EGO-1, Dicer-related helicase DRH-3 and Tudor domain protein EKL-1 (Claycomb et al., 2009), then uridylated by nucleotidyltransferase CDE-1 to restrict the association of these 22G-siRNA to argonaute CSR-1 only (van Wolfswinkel et al., 2009).

Instead of simply silencing its targets, whole genome microarray analysis revealed the complex effects of CSR-1 depletion (Claycomb et al., 2009). The pathway has been implicated in the fine tuning of germline expression, mRNA maturation, protecting targets against piRNA-induced silencing, and translational regulation (Avgousti et al., 2012, Friend et al., 2012, Seth et al., 2013, Wedeles et al., 2013b, Gerson-Gurwitz et al., 2016). CSR-1 RNAi pathway is the only RNAi pathway required for proper chromosome segregation in *C. elegans*, but its exact role in chromosome segregation remains elusive (Yigit et al., 2006, Claycomb et al., 2009). Gerson-Gurwitz et al. suggested the chromosome missegregation in *csr-1* mutant is mainly caused by misregulation of multiple embryonic cell division genes. In particular, overexpression of KLP-7, a kinesin-13 microtubule depolymerase, in *csr-1* mutant accounts for its microtubule assembly defect, but not the chromosome segregation defect (Gerson-Gurwitz et al., 2016). 89% of CSR-1-targeted germline-expressing genes have low HCP-3 occupancy (Gassmann et al., 2012). We hypothesized that CSR-1 affects centromere protein HCP-3 occupancy, and the chromosome missegregation in CSR-1 depletion is a result of defective centromere function. The overloading of centromeric and kinetochore proteins on the mitotic chromosomes may cause chromosome-microtubule misattachments. Specifically, if the sister kinetochore attaches to microtubules emanated from the two poles, it will lead to merotelic attachments and lagging chromosomes (Cimini et al., 2001).

Here, we have characterized mitotic defects related to the defective centromere function and kinetochore attachment in the CSR-1-depleted embryos. Cell division timing, spindle pole separation dynamics, and congression of chromosomes were examined. We also evaluated how CSR-1 knockdown affects HCP-3 expression, organization, and the assembling of a functional holocentromere and kinetochore. While the total centromere protein HCP-3 level was not increased, its level on the chromosome increased in CSR-1 knockdown. We further deciphered which CSR-1 isoform and domain influences HCP-3, and whether the change of HCP-3 on chromatin depends on the RNAi pathways and HCP-3 loading pathway.

## Results

### CSR-1 knockdown embryos are prone to erroneous chromosome segregation

Chromosome missegregation and aneuploidy have been reported in *csr-1* hypomorph mutant and RNAi knockdown embryos through immunofluorescence and fluorescence *in situ* hybridization (Claycomb et al., 2009). Our RNAi treatment reduced *csr-1* mRNA to 16% of its original level (**Figure S1A**) and results in over 80% embryonic lethality (**Figure S1B**). 70 % of *csr-1*(RNAi) 1-cell embryos exhibited chromosome missegregation with lagging chromosomes (**Figure 1A**) and 30% *csr-1*(RNAi) embryonic cells were aneuploid for the marked chromosome (**Figure 1B**). We also noted a weaker α–tubulin signal in the *csr-1*(RNAi) embryos by immunofluorescence (**Figure 1A**).

**Figure 1.**
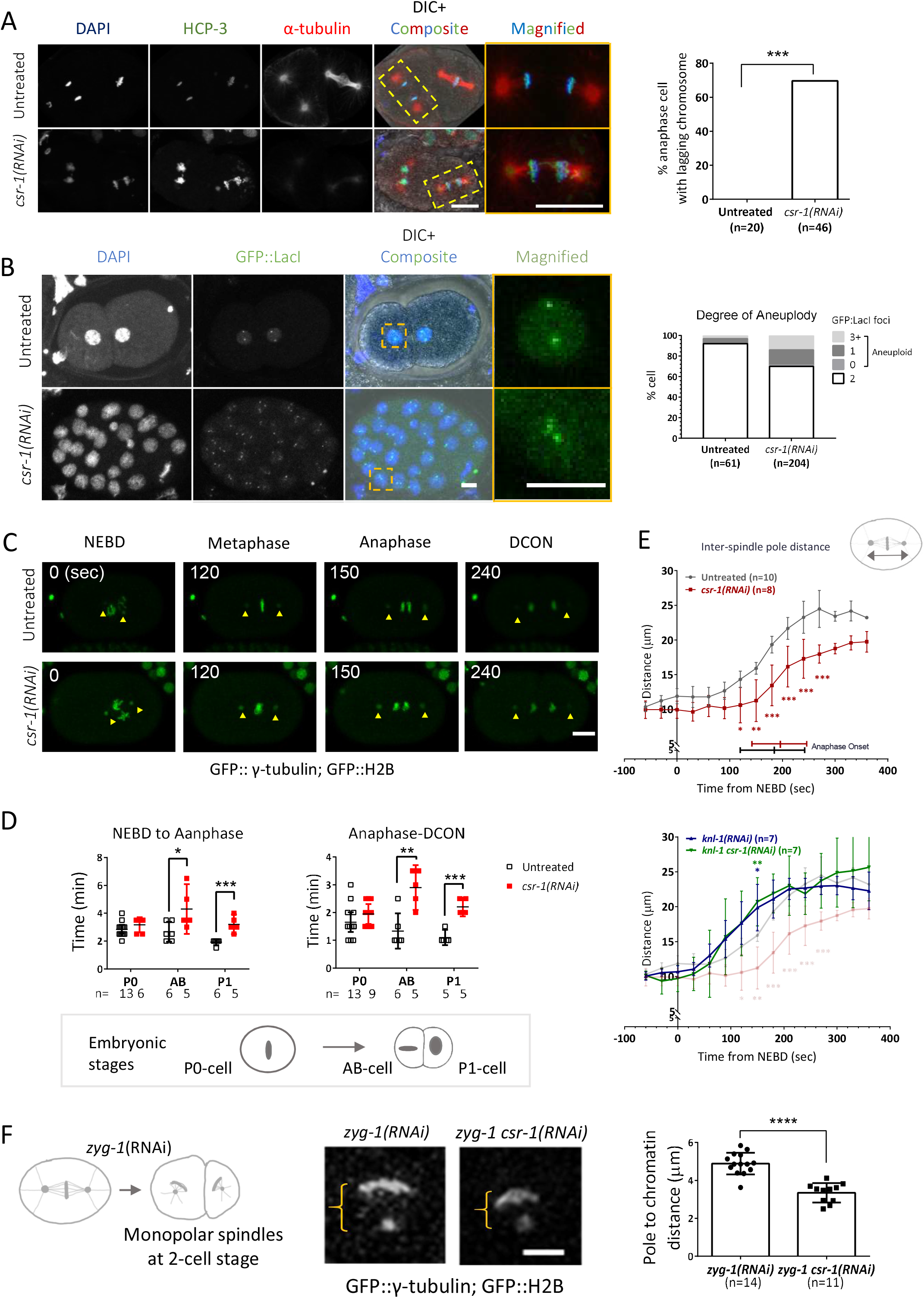
CSR-1 depletion affects chromosome segregation. (A) Untreated control or *csr-1*(RNAi) embryos are immunostained for HCP-3 and α-tubulin with DAPI-staining. Examples of a normal versus a missegregated anaphase chromosomes are highlighted in dashed rectangles and enlarged on the right. Scale bars: 10 μm. Percentage of lagging chromosomes observed were quantified. n equals the number of anaphase cells analyzed. ***: p<0.001, Fisher’s exact test. (B) Embryos containing GFP::LacI-labelled LacO repeats on the translocated Chromosome II/III were immunostained for GFP with DAPI staining, showing the copy number of the translocated chromosome. Examples of a normal cell versus an aneuploid cell are highlighted in dashed rectangles and enlarged on the right. Scale bars: 5 μm. Percentage of aneuploid cells were scored in the untreated and *csr-1*(RNAi) embryos. n equals the number of cells analyzed. ***: p<0.001, fisher’s exact test. (C) The spindle poles (yellow arrows) and chromatin are tracked in the 1^st^ embryonic division in a strain expressing GFP::γ-tubulin and GFP::H2B. Time from NEBD are indicated on top left. Scale bar: 10 μm. NEBD: nuclear envelope breakdown; DCON: DNA condensation. (D) Time spent in early and late mitotic phases of the 1-cell embryos (P0) and the 2-cell embryos (AB and P1) quantified from live-cell imaging as shown in (C). n equals the number of embryos analyzed. (E) Distance between spindle poles in mitosis in one-cell embryos with different RNAi treatments. Top: untreated control and *csr-1*(RNAi) embryos. Bottom: Kinetochore protein KNL-1 knockdown (MT-KT detachment) with or without csr-1(RNAi) co-depletion. n equals the number of embryos analyzed. The timing for anaphase onset for untreated and *csr-1*(RNAi) P0 embryos are extracted from (D). (F) zyg-1(RNAi) inhibits centriole duplication and generates monopolar spindle atat 2-cell stage. Live-cell imaging shows the congressed chromatin mass around the sole spindle pole in cells expressing GFP::γ-tubulin and GFP::H2B with or without *csr-1*(RNAi) co-depletion. The distance between the spindle pole and the congressed chromatin was quantified. Scale bar: 5 μm. n equals the number of monopolar spindle cells analyzed.

### CSR-1 knockdown prolongs mitosis in 2-cell stage

The mitotic time spent in the first two cell divisions were quantified by live-cell imaging. A worm strain expressing mCherry::H2B was used for marking the mitotic events **(Figure 1C)**. Time from prophase to anaphase and time from anaphase to telophase were taken as early and late mitosis, respectively **(Figure 1D)**. In the first embryonic division the embryonic founder cell (P0) showed no significant difference in mitosis timing. In the 2-cell embryos, AB and P1 cells exhibited prolonged early mitosis (1.6-fold and 1.7-fold) and late mitosis (2.2-fold and 2-fold) when compared to the untreated control. Delayed anaphase onset can occur when the spindle checkpoint is activated (Essex et al., 2009). The lengthened late mitosis suggests the possibility that the chromosomes are oriented in a way that requires more time to separate, for example if they are tangled or misattached to the microtubules (Gregan et al., 2011), or alternatively, there could be a weaker astral pulling force to pull the chromosomes apart (Grill et al., 2001).

### CSR-1 knockdown slows down spindle pole separation which is kinetochore-dependent

Chromosome-to-pole movement is minimal during chromosome segregation in *C. elegans* as the process is predominantly driven by the shortening of astral spindles (Oegema et al., 2001). To measure spindle pole separation in mitosis, the distance between the two spindle poles in 1-cell embryos expressing GFP::⍰-tubulin was measured. Knockdown of CSR-1 significantly delayed the onset and reduced the extent of pole separation **(Figure 1E)**. The terminal distance between spindle poles in *csr-1*(RNAi) embryos was reduced by 9% at the mitotic exit. To determine if the delay is kinetochore-dependent, outer kinetochore protein KNL-1 was depleted singly or together with CSR-1. knl-1(RNAi) embryos had premature pole separation, while knl-1 *csr-1*(RNAi) phenocopied the knl-1(RNAi) **(Figure 1E)**. Thus, the observed pole separation delay in *csr-1*(RNAi) was dependent on the presence of KNL-1, which implies the dependence on kinetochore-microtubule attachment. On the other hand, the astral spindle pulling force which is kinetochore-independent is normal in *csr-1*(RNAi). Aurora B kinase AIR-2 helps resolve misattached kinetochore-microtubule to allow re-connection (Kaitna et al., 2002), but the localization of AIR-2 was not altered in CSR-1 knockdown embryos **(Figure S1C)**.

### CSR-1 knockdown affects chromosome position

To generate a scenario without any merotelic attachments and observe the relative position of chromosome to the spindle pole, we observe the situation with only one spindle pole. ZYG-1 is a centrosome protein important for centrosome duplication (Caron et al., 1999). Without ZYG-1, the centrosome fails to replicate which gives rise to cells with only one spindle pole after the first division, so this excluded any merotelic attachment (Powers et al., 2004). Congressed chromosomes in these cells form an arc-shape around the spindle pole at late prometaphase **(Figure 1F)**. Measurement of the relative position of the chromatin to the spindle pole reflects the balance between kinetochore pulling force toward the pole and kinetochore-independent pole repulsion force (Bajer and Molè-Bajer, 1972, Dumont et al., 2010). In *zyg-1*(RNAi), the average distance between the spindle pole and the center of mass of the chromatin was measured to be 4.891 μm. *zyg-1 csr-1*(RNAi) cells have a shorter average chromatin-to-pole distance of 3.349 μm **(Figure 1F)**. Thus, the abnormality in chromosome positioning in *csr-1*(RNAi) cells exists even if there is no merotelic attachment.

### CSR-1 knockdown alters some, but not all, kinetochore protein levels on mitotic chromosomes

Next, kinetochore protein recruitment to mitotic chromosomes was evaluated. Fluorescent labeled kinetochore proteins were tracked in metaphase plate by live cell imaging **(Figure 2A)**. The total intensities of HCP-3, MIS-12, KNL-1, KNL-3, and HCP-1 increased significantly in *csr-1*(RNAi) embryos whereas histone H2B intensity was not significantly different. The metaphase plates appear bulkier, which is consistent to the chromosome misalignment phenotype previously reported in *csr-1*(RNAi) embryos (Claycomb et al., 2009, Gerson-Gurwitz et al., 2016). The imperfectly aligned mitotic chromosomes, thus increased volume of the chromosomes, cannot explain all about the increase in the total intensity in *csr-1*(RNAi) embryos. The average HCP-3 signal on metaphase chromatin also increased in *csr-1*(RNAi) embryos **(Figure S2A)**.

**Figure 2.**
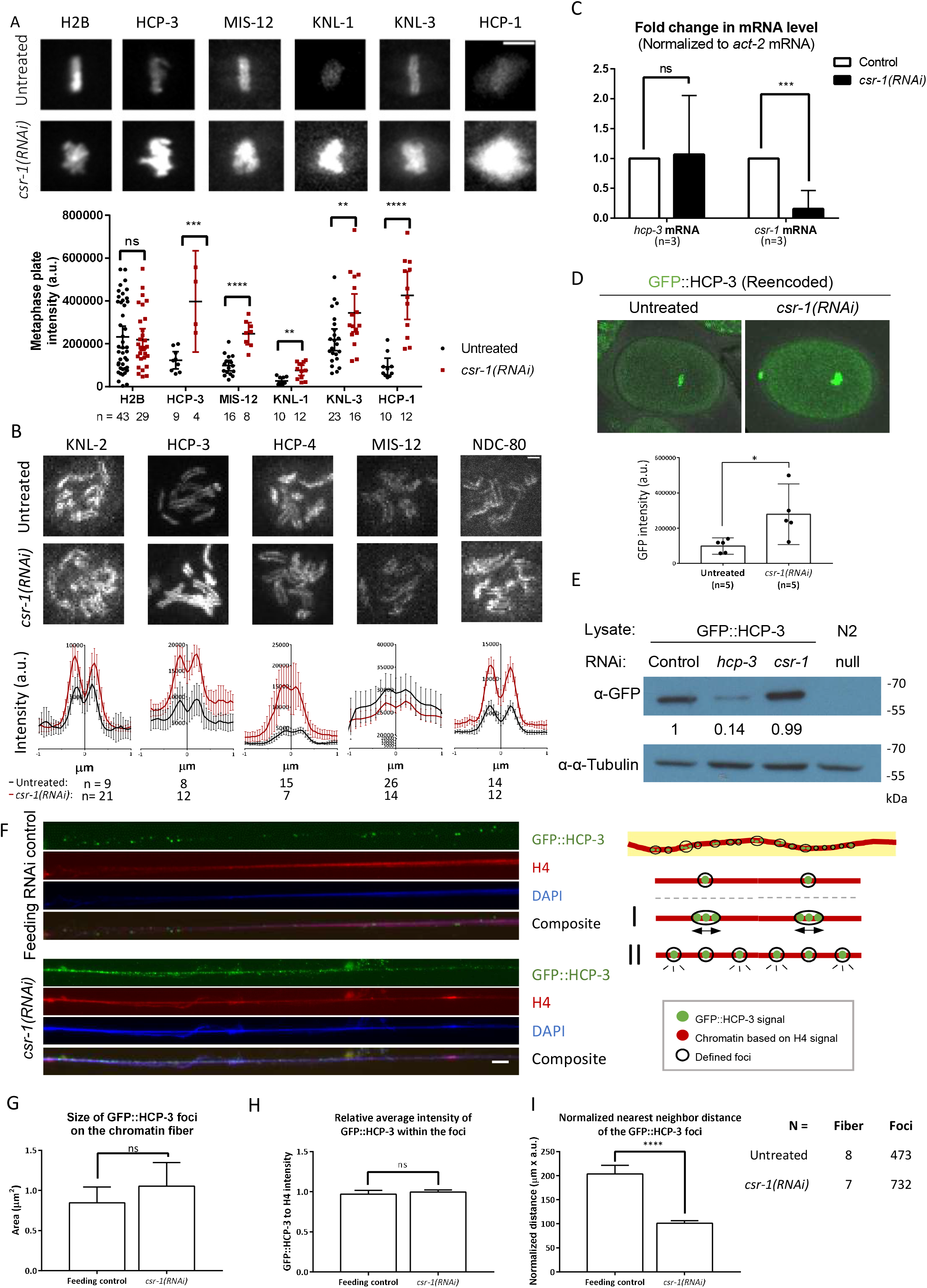
CSR-1 depletion affects kinetochore and centromere protein localization. (A) Maximum-projected live-cell images of metaphase chromosomes in the untreated and the *csr-1*(RNAi) embryos. Total metaphase fluorescent intensities in 1-cell embryos expressing different kinetochore proteins fused with fluorescent protein were quantified. n equals the number of embryos analyzed. Scale bar: 5 μm. (B) Immunofluorescence image of prometaphase chromosomes from a single plane in the untreated and the *csr-1*(RNAi) embryos. Scale bar: 1 μm. Either 1 or 2-cell embryos were used. Mid-point between two peaks was taken as the center (0 μm) to align the linescan intensity profiles of different chromosomes. n equals the number of chromosomes analyzed. (C) Relative fold change of hcp-3 and *csr-1* mRNA levels in the *csr-1*(RNAi) embryos compared to the feeding RNAi control. The mRNA levels are normalized to that of act-2. Three biological replicates were performed. (D) Knockdown CSR-1 in strain with hcp-3 gene reencoded. The strain contains reencoded hcp-3 gene from OD1174 with endogenous hcp-3 knockout allele from VC1393. GFP intensity of the metaphase plate was quantified. n equals the number of embryos analyzed. Scale bar: 10 μm. (E) Immunoblot shows that GFP::HCP-3 protein level remains unchanged in the *csr-1*(RNAi) embryos when compared to the feeding RNAi control. Loading control: α-tubulin. Ladder sizes are indicated on the right. N2: wildtype lysate, biological control without GFP::HCP-3. Numbers below α-HCP-3 blot indicate the normalized fold-change of HCP-3 level from 3 biological replicates (Figure S2B). (F) Representative images of the chromatin fibers prepared from control feeding RNAi and *csr-1*(RNAi). Scale bar: 10 μm. Right: diagram explaining how the GFP::HCP-3 foci are defined. 20. 20-pixel-wide line (yellow background) were drawn across the length of fiber (red). Below are illustrations showing 2 scenarios of HCP-3 mislocalization on the fibers, I: increase in foci size and/or intensity; II: reduction in nearest neighbor distance. A GFP-intensity threshold was applied (see Figure S2D) to define the GFP::HCP-3 foci for quantifications shown in (G-I). (G-I) Properties of the GFP::HCP-3 foci are quantified, including the foci size, foci average intensity, and foci density. The GFP::HCP-3 foci intensities were normalized with corresponding H4 intensities. Foci density was assessed by nearest neighboring distance (NND) for each focus and normalized to the inverse of corresponding H4 intensity. RNAi-control: n= 473 foci from 8 fibers; *csr-1*(RNAi): n= 723 foci from 7 fibers.

To examine the localization of different kinetochore proteins on individual chromosomes, kinetochore proteins levels were immune-stained and imaged with higher resolution. Average intensities of a 10-pixel-width line along the lateral axis of individual prometaphase chromosomes were profiled **(Figure 2B)**. As reported in Gerson-Gurwitz et al. (2016), the centromere structure remains bipolar in *csr-1*(RNAi) embryos. While the M18BP1 homolog KNL-2 and kinetochore protein MIS-12 levels were not significantly altered, the levels of the kinetochore proteins HCP-3, HCP-4 and NDC-80 on the prometaphase poleward faces were all elevated in *csr-1*(RNAi) embryos. HCP-3 is on the top of the localization hierarchy among these proteins (Oegema et al., 2001). Our results suggested that at least some of the kinetochore proteins increased their localization at the holocentromere due to CSR-1 depletion. Note that MIS-12 level was found to be increased by live-cell imaging during metaphase but remained unchanged in prometaphase by immunofluorescence. GFP-tagged MIS-12 were quantified in the live imaging assay whereas MIS-12 antibody was used in immunofluorescence for untagged MIS-12. Yet, the reason for the discrepancy is not clear.

### CSR-1 knockdown does not affect HCP-3 mRNA nor protein levels

CENP-A overexpression in other organisms usually leads to excess CENP-A loading, which induces formation of ectopic centromeres (Tomonaga et al., 2003, Heun et al., 2006, Amato et al., 2009). To decipher how CSR-1 modulates the HCP-3 level on the chromatin, we tested if CSR-1 knockdown increases HCP-3 mRNA or protein level. The strain contains a sole copy of HCP-3 that is GFP-fused by MosSCI in a hcp-3 deletion background hcp-3(ok1892) (Gassmann et al., 2012). Target genes were knocked down by feeding RNAi to collect sufficient amount for RT-qPCR and Western blot. In *csr-1*(RNAi) embryos, hcp-3 mRNA level was not significantly different from the feeding RNAi control **(Figure 2C)**. Similarly, the total embryonic GFP::HCP-3 protein level of the *csr-1*(RNAi) was not significantly different from the feeding RNAi control **(Figure 2D)**.

We also tested whether the CSR-1 repression depends on specific hcp-3 endogenous sequence. However, with hcp-3 gene reencoded (Gerson-Gurwitz et al., 2016), the GFP::HCP-3 level still increased upon the depletion of CSR-1, which is 2.8-fold of the untreated control **(Figure 3F)**. Additional HCP-3 loading does not seem to depend on specific, endogenous hcp-3 mRNA sequences.

**Figure 3.**
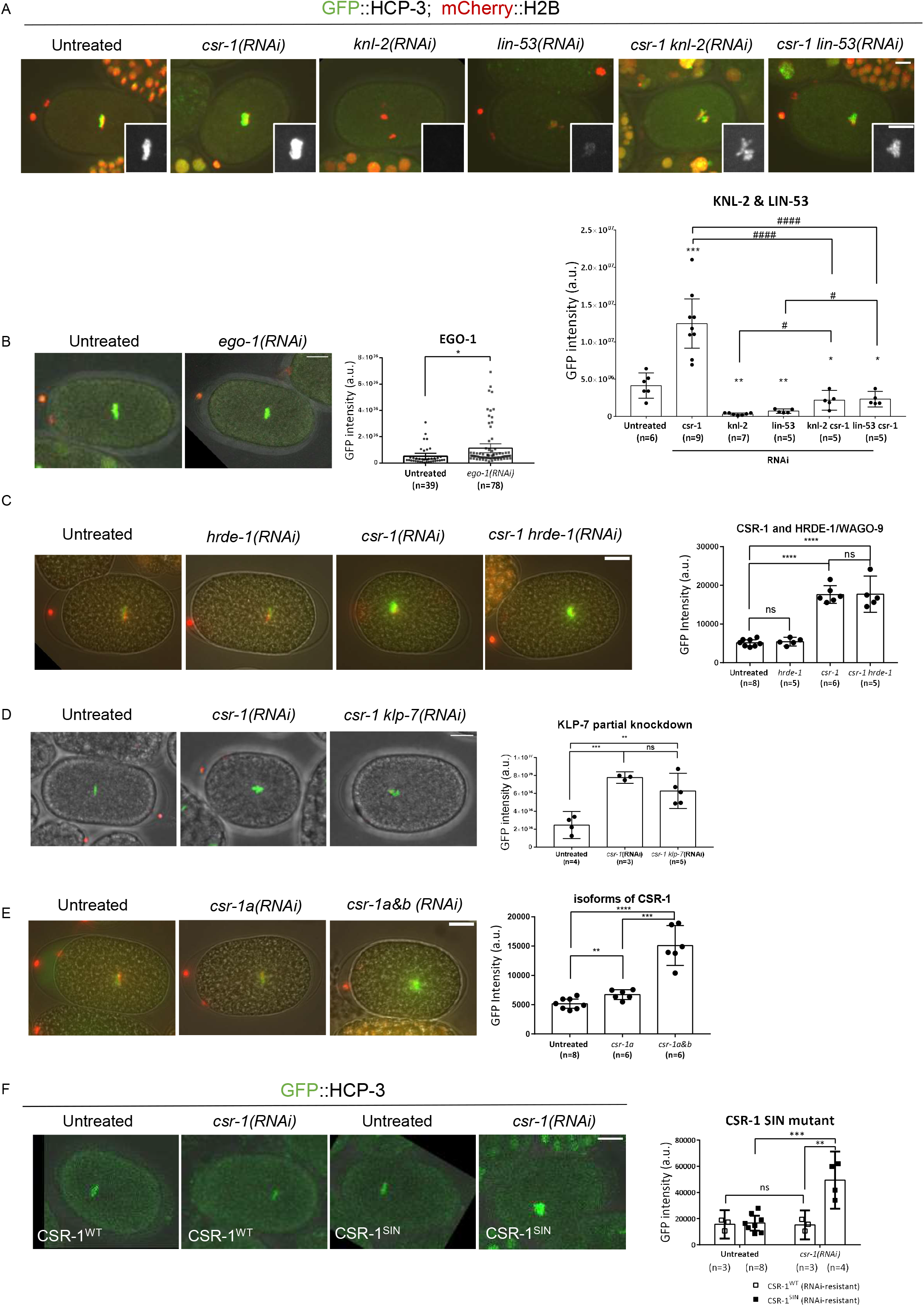
Requirement of CSR-1’s repression of HCP-3 chromatin localization. Quantification of the GFP::HCP-3 intensity of the metaphase plate in one-cell embryos with different RNAi treatments. (A) Knockdown of HCP-3 loading factors: KNL-2 and LIN-53. Insets showing the metaphase GFP signal with higher contrast. (B) Knockdown of the RdRP of CSR-1 RNAi pathway: EGO-1. (C) Knockdown of the argonaute HRDE-1 of another nuclear RNAi pathway. (D) Partial knockdown of KLP-7 in *csr-1*(RNAi) embryos. n equals number of one-cell embryos quantified. (E) Knockdown of specific CSR-1 isoform A alone. (F) Knockdown CSR-1 in strains expressing transgenic, RNAi-resistant CSR-1 (wildtype or SIN, slicer inactive mutant). *: comparison to the untreated; #: comparison between RNAi groups. Scale bar: 10 μm.

### CSR-1 knockdown increases chromatin HCP-3 level on mitotic chromosomes

The turnover of HCP-3 on chromosome was assayed by photobleaching assay as described in Gassmann et al. (2012). GFP::HCP-3 on one of the anaphase sister in 1-cell stage was photobleached with 488nm laser. The GFP signal on the next metaphase represents the amount of new GFP::HCP-3 loaded within one cell cycle. The GFP signals of the two daughter cells, one bleached and one unbleached, were quantified. The ratio of GFP on the bleached to unbleached (B:U) chromosomes did not drop in csr-1(RNAi) embryos when compared to that in the untreated control **(Figure S2C)**.

To evaluate HCP-3 distribution on the *csr-1*(RNAi) chromosomes, chromatin were stretched into fibers based on Kyriacou and Heun (2018). Consistent with the in vivo imaging, *csr-1*(RNAi) chromatin fibers contain more GFP::HCP-3 staining **(Figure 2F)**. We defined GFP foci on the chromatin fiber by manual thresholding and dots clustering. The area and the mean focal intensity of the GFP foci on *csr-1*(RNAi) fibers were not significantly different from the feeding control (Figure 2G & H). The density of the defined GFP foci was measured by the nearest neighboring distance (NND). The normalized NND of GFP foci on the *csr-1*(RNAi) fibers was significantly lower than that on the feeding RNAi control fibers (Figure 2I).

### Some HCP-3 chromatin localization are independent on KNL-2 in *csr-1*(RNAi) embryos

Centromere licensing factor Mis18BP1/KNL-2 and histone chaperone RbAp46/48/LIN-53 are the two known factors required for HCP-3 localization (Maddox et al., 2007, Lee et al., 2016). To determine whether the loading requirement of the HCP-3 in *csr-1*(RNAi) embryos is different, KNL-2 or LIN-53 were co-depleted with CSR-1. Co-depleting CSR-1 and KNL-2 gave rise to dot-like chromatin and severe anaphase bridging as in knl-2(RNAi) embryos **(Figure 3A)**. While GFP::HCP-3 signal almost abolished on chromatin in knl-2(RNAi) embryos, noticeable GFP::HCP-3 were observed on the dot-like chromatin mass in *csr-1* knl-2(RNAi) embryos, whose intensity equals 6.5-fold of the knl-2(RNAi) and 53% of the untreated control **(Figure 3A)**. Co-depletion of CSR-1 and LIN-53 gave rise to lagging chromosome as in lin-53(RNAi) embryos, and the GFP intensity shown a 3-fold increase compared to lin-53(RNAi), and 56% of the untreated control **(Figure 3A)**. These results indicated that in *csr-1*(RNAi) embryos, at least a proportion of the HCP-3 can localize onto the chromatin independently of KNL-2 or LIN-53.

### The HCP-3 is restricted via CSR-1 RNAi pathway, but is not dependent on the antagonized HRDE-1 RNAi pathway

EGO-1 is the RNA-dependent RNA polymerase in CSR-1 RNAi pathway important for the biogenesis of the 22G-siRNA. HRDE-1 is the argonaute of another nuclear RNAi pathway, in which CSR-1 is proposed to protect endogenous germline genes from silencing by HRDE-1 (Seth et al., 2013, Xu et al., 2018). To determine if the repression of HCP-3 loading functions via RNAi pathways, EGO-1 or HRDE-1 was depleted **(Figure S3A)**. GFP intensity in the ego-1(RNAi) embryos is 2-fold of the untreated embryos **(Figure 3B)**.

If the CSR-1-protection of germline expression against HRDE-1 silencing is coupled to the restriction of HCP-3 localization, then after co-depletion, we would expect to see HCP-3 level back to the untreated level. As HRDE-1 depletion is not embryonic lethal, the knockdown efficiency was verified by RT-qPCR **(Figure S3B)**. Co-depletion of HRDE-1 and CSR-1 did not prevent excess HCP-3 chromatin localization. The GFP intensities in both *csr-1*(RNAi) and *csr-1* hrde-1(RNAi) are 3.4-fold of the untreated control **(Figure 3C)**. Thus, CSR-1-repressed HCP-3 localization does not depend on HRDE-1.

### The increase in HCP-3 chromatin localization was not dependent on CSR-1’s target KLP-7 overexpression

To investigate if CSR-1’s effect on HCP-3 localization is indirect through the misregulation of specific CSR-1 targets, we tried to rescue the phenotype by reversing the level of a specific CSR-1 target, KLP-7. We partially depleted KLP-7 together with CSR-1. The co-depletion alleviated the alignment defect at metaphase but does not improve the chromosome segregation accuracy as reported previously **(Figure S3C)**. The quantification of metaphase HCP-3 level is similar for the embryos with or without KLP-7 co-depletion **(Figure 3D)**. Hence, the elevation of HCP-3 level is independent of KLP-7 upregulation.

### The increase in HCP-3 chromatin localization requires both CSR-1 isoforms

CSR-1 have two isoforms, which have different localizations and were proposed to have different functions (Avgousti et al., 2012). The larger isoform CSR-1A is more abundant in soma cells whereas CSR-1B is more abundant in germlines (Claycomb et al., 2009). The two isoform genes differ by one exon at the N-terminus. In this study, we use dsRNA targeting the common sequence of both isoforms for *csr-1*(RNAi) **(Figure S3D)**. To deplete only CSR-1A, we use dsRNA targeting the first exon of isoform A. As depletion of CSR-1A alone is not embryonic lethal (Gerson-Gurwitz et al., 2016, Charlesworth et al., 2021), we validated the *csr-1*a knockdown by RT-PCR, whose mRNA level is 12.23% of that in the untreated embryos **(Figure S3B)**. Depletion of CSR-1A results in 1.3-fold increase in chromatin GFP::HCP-3 level in 1-cell stage embryos, compared to the untreated control **(Figure 3E)**. However, the increase of the GFP::HCP-3 level in CSR-1A depletion is not as much as in CSR-1A and B depletion, which is 3-fold of the untreated control. Moreover, depletion of CSR-1A does not affect chromosome alignment and chromosome segregation in early embryos **(Figure S3C)**.

### The increase in HCP-3 chromatin localization was dependent on the PIWI domain activity

‘DDH’ motif in the PIWI domain of CSR-1 is important for mRNA cleavage by binding the divalent magnesium ions (Liu et al., 2004, Schwarz et al., 2004). CSR-1 cuts target mRNA in vitro (Aoki et al., 2007). A CSR-1 slicing inactive mutant (CSR-1^SIN^) with point mutations D606A and D681A has been shown to induce chromosome missegregation in the absence of the endogenous CSR-1 (Yigit et al., 2006, Aoki et al., 2007, Gerson-Gurwitz et al., 2016). We examined GFP::HCP-3 metaphase intensity in strains expressing the reencoded, hence RNAi-resistant CSR-1^WT^, and CSR-1^SIN^ (Figure 3F). After endogenous CSR-1 RNAi is knocked down, embryos expressing CSR-1^SIN^ showed a significant increase in the GFP::HCP-3 intensity, when compared to embryos expressing CSR-1^WT^. Thus, CSR-1-repressed HCP-3 localization requires CSR-1’s PIWI domain slicing activity.

## Discussion

We investigated the role of CSR-1 in chromosome segregation by analyzing microtubules and kinetochore function in multiple embryonic cell cycle events **(Figure 4A)**. The chromosome segregation at anaphase is driven by the shortening of astral spindles. Astral spindles connect spindle poles to the cell membrane, and mitotic spindles connect spindle poles to the chromosomes. Although spindle poles separation is slower in *csr-1*(RNAi) embryos, CSR-1 depletion does not affect the astral spindle depolymerization. The spindle pole separation speeds up when they are detached from the kinetochore in knl-1 *csr-1*(RNAi) embryos. Thus, defective spindle pole separation in *csr-1*(RNAi) is dependent on spindle-kinetochore attachment. The attachment is likely merotelic, a situation where one kinetochore connects to both spindle poles. Merotelic attachment can also hinder chromosome alignment at metaphase. We tested if eliminating possible merotelic attachment prevents difference in the way chromosomes position. In *zyg-1 csr-1*(RNAi) embryonic cells with monopolar spindles, the chromosome-pole distance becomes shorter. This observation implied a change in the net force acting on the chromosomes, by either a stronger pole attraction force or a weaker pole repulsion force. Changes in pole repulsion force can affect chromosome movements and their alignment at the equator at metaphase (Antonio et al., 2000, Guo et al., 2013), which could in turn hinder amphitelic kinetochore-microtubule attachment (Powers et al., 2004). Weaker pole repulsion force can result from excessive microtubule depolymerization, which have been previously observed in *csr-1*(RNAi) embryos (J. Richard McIntosh et al., 2002). KLP-7, the homolog of mitotic centromere-associated kinesin (MCAK) in *C. elegans*, whose expression is suppressed by CSR-1 (Claycomb et al., 2009), limits microtubule outgrowth from centrosomes (Srayko et al., 2005). Reducing KLP-7 level in csr-1(RNAi) embryos rescued the microtubule assembly defect and improved the alignment of chromosomes at metaphase plates (Gerson-Gurwitz et al., 2016). However, in these embryos, chromosomes still missegregate, which may arise from defective centromeres or kinetochore functions.

**Figure 4.**
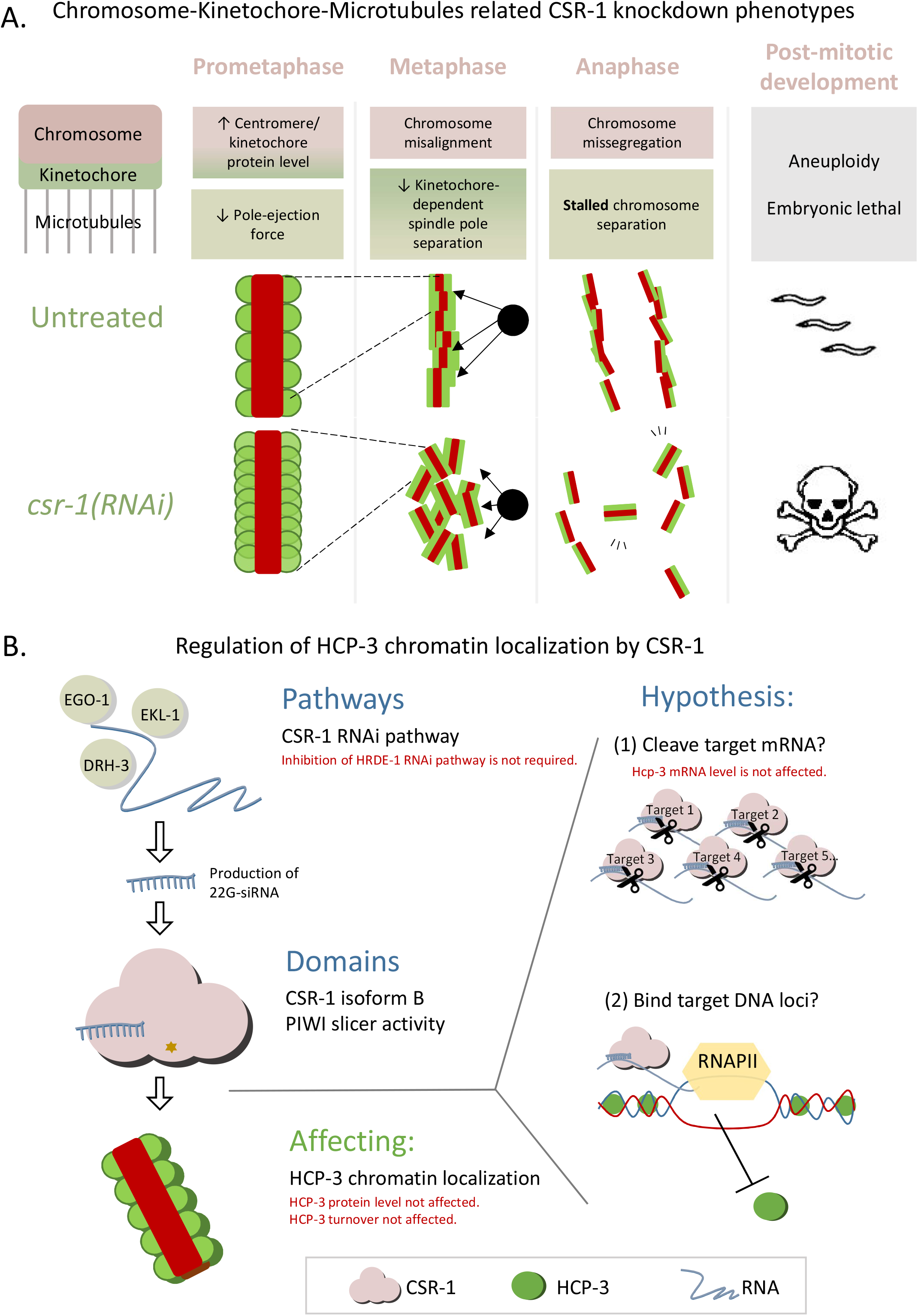
The current understanding of CSR-1 RNAi in regulating HCP-3 localization on centromere. (A) A summary of CSR-1 knockdown phenotypes related to chromosome segregation in mitosis. (B) A proposed mechanism for how CSR-1 affects HCP-3 centromere localization. The germline-enriched CSR-1 isoform B with its PIWI domain is important for regulating HCP-3’s loading onto chromatin. This could be achieved through regulating (1) the level of a target or a subset of mRNA targets by cleaving mRNA as proposed in Gerson-Gurwitz et al. (2016) or (2) an epigenetic mark inhibitory for HCP-3 loading on the target genome via physical interaction in RNAPII-dependent manner as proposed in Wedeles et al. (2013b). Pathways and CSR-1 domains found to be important for HCP-3 repression are summarized in black. Factors that we have tested to be not required or negative are shown in red.

Among the proteins with elevated levels, HCP-3 is on the top of the kinetochore protein recruitment hierarchy. Centromeric protein HCP-3 on prometaphase chromosomes remained bipolar in our linescan analysis **(Figure 2F)**, as in Gerson-Gurwitz et al. (2016). Yet, more HCP-3 was binding onto the poleward faces on the chromosomes, suggesting that CSR-1 suppressed HCP-3 localization on centromere. None of the HCP-3 mRNA, protein, nor its chromatin turnover rate changed after CSR-1 depletion **(Figure 2C-E)**. With the total level unchanged in embryos, more HCP-3 protein localized to the mitotic chromosomes when CSR-1 was knocked down. It implies that in the untreated condition, not all embryonic HCP-3 are incorporated to the chromatin, and its chromatin level is normally suppressed by CSR-1. Recruitment of HCP-3 onto the holocentromere is interdependent on KNL-2 and partially dependent on LIN-53 in embryos (Maddox et al., 2007, Lee et al., 2016). When CSR-1 is depleted, such dependency no longer holds true, as some level of KNL-2- and LIN-53-independent HCP-3 localization was observed (Figure 3A). This phenomenon is interesting and it hints that part of the function of CSR-1 is to suppress HCP-3’s non-specific chromatin localization. Reducing the HCP-3 level does not rescue the chromosome missegregation phenotype in the CSR-1 and partial HCP-3 co-depletion conditions we tested **(Figure S3E)**. We propose that the organization pattern of HCP-3 is altered in *csr-1*(RNAi) embryos, as HCP-3 foci density on stretched chromatin fibers increases.

Ladouceur et al. (2017) have examined the localization of HCP-3 on the squashed prometaphase chromatin. HCP-3 dots have similar size and distribution, but HCP-3 dot intensity is higher after *csr-1*(RNAi) treatment. By stretching chromatin into fibers, we observed HCP-3 localization as foci of similar size and brightness, but with increased density after *csr-1*(RNAi) treatment. Potentially, our chromatin fibers resolved HCP-3 to a higher degree, so within one brighter HCP-3 dot in the squashed chromatin, there could be multiple HCP-3 foci. If the HCP-3 foci in our chromatin fibers represent part of the broader HCP-3-enriched domain as observed in chIP-chip or chIP-seq studies (Gassmann et al., 2012, Steiner and Henikoff, 2014), our result of a reduced distance between HCP-3 foci suggests that HCP-3 nucleosomes may become more dense within the HCP-3 domain (Graphic abstract). We noticed that there were more HCP-3-containing fibers in *csr-1*(RNAi) than in control as we selected for GFP positive chromatin fibers. We therefore cannot exclude the possibility that HCP-3 domains expanded into non-permissive sites. However, either denser or more HCP-3 domains can provide more places for more kinetochore proteins to bind, as this is what we observed for HCP-4 and NDC-80. Additional microtubule binding sites may nucleate on chromosomes.

The nature of CSR-1’s suppression on HCP-3 remains unclear. CSR-1 may suppress HCP-3 either through indirect transcript regulation or direct genome interference **(Figure 4B)**. Candidate CSR-1 targets that affect HCP-3 loading include histone chaperones or other histones. CSR-1 may suppress HCP-3 localization by limiting the expression of histone chaperones that place HCP-3 onto the chromosomes. However, expression levels of LIN-53 and KNL-2 remain normal in CSR-1-depleted embryos or in CSR-1 SIN mutant (Claycomb et al., 2009, Gerson-Gurwitz et al., 2016). Moreover, CSR-1 slices the pre-mRNA of canonical histone mRNAs to promote its maturation (Avgousti et al., 2012). This may affect the HCP-3 incorporation by altering the level of mature H3, H4 mRNAs, as the H3:H4 ratio has been shown to affect CENP-A^Cnp-1^ incorporation in fission yeast (Castillo et al., 2007).

Another emerging proposal we favor is that CSR-1 may directly regulate epigenetic marks on genome. CSR-1 can localize onto its targets’ genomic loci, likely by binding to nascent mRNA target as it co-immunoprecipitates with RNAPII (Claycomb et al., 2009, Wedeles et al., 2013b, Tu et al., 2015). CSR-1-targeted gene loci were shown to be enriched with certain histone modifications by chIP, such as H3K4me2,3 and H3K36me2,3 (Gerstein et al., 2010, Wedeles et al., 2013a). CSR-1 has been shown to exhibit transgenerational protection of its targets from piRNA-mediated silencing, likely via creating protective epigenetic marks on its targets’ gene loci (Wedeles et al., 2013b). It will be interesting and important to explore the contribution of epigenetic modifications including histone marks and CSR-1-genome interaction on restricting HCP-3 localization. However, HRDE-1, one of the argonautes essential for maintaining the gene silencing status, is not required for CSR-1’s ability to suppress HCP-3. Thus our results indicated that the suppression does not rely on CSR-1’s antagonistic functions with the HRDE-1 RNAi pathway. The role of CSR-1’s PIWI domain in HCP-3 suppression is elusive in this hypothesis. While the 2 mutations at the PIWI domain are crucial for its slicing activity, it is possible that they are required for CSR-1 binding with the other cofactors to suppress HCP-3, for example glycine-tryptophan (GW) repeat proteins binds to argonaute at the PIWI domain for translational repression (Liu et al., 2005, Eulalio et al., 2008). The PIWI domain SIN mutant has recently been shown to change the 22G-siRNA abundance and its profile (Singh et al., 2021). This implied that how the argonaute interacts with the target mRNAs, and with which mRNAs as targets could also be affected in the CSR-1^SIN^ mutant.

In this study, we have identified CSR-1 as a repressor against excess HCP-3 centromere localization in *C. elegans*. CSR-1 suppresses HCP-3 via its RNAi pathway and with its slicing activity. The regulation likely happens in the hermaphrodite germline where the CSR-1 isoform b is enriched (Charlesworth et al., 2021). Specific depletion of the other soma cells-enriched isoform a does not cause embryonic lethality (Claycomb et al., 2009, Gerson-Gurwitz et al., 2016), chromosome missegregation, as well as a dramatic GFP::HCP-3 elevation as the depletion of both isoforms does. Our study has shed light on how holocentromeric regions are distinguished from the other chromatin regions in *C. elegans*, and exemplified how RNAi pathway can be involved in the determination of centromere regions in organisms containing non-regional centromeres. In humans, the AGO-2 RNAi pathway is also required for proper chromosome segregation. Depleting of AGO-2, or with Slicer-inactive AGO-2 mutant, leads to abnormal level of α-satellite RNA and CENP-C1 mislocalization to the chromosome arms (Huang et al., 2015). It will be interesting to investigate if argonautes and RNAi pathways have conserved function for maintaining centromere or controlling CENP-A loading. Even though humans and *C. elegans* have different centromere architectures and DNA sequences, the epigenetic regulation, including histone modifications, RNAi pathways, transcription and non-coding RNAs, may follow general rules.

## Acknowledgement

We thank Yuen lab members for suggestions, Kyle Ka Lun Law for his pioneer trials, Lok Yee Hiok for generating the worm strain and Yick Hin Ling for discussion. We thank Karen Oegema and Arshad Desai lab for providing antibodies and strains, Julie Claycomb lab for providing strain and discussion, Garry Wong lab for providing the feeding RNAi plasmid and suggestions, and E. Kyriacou and Patrick Heun for suggestions on chromatin fiber assay. This work was supported by grants General Research Funds 17126717 and 17116520 by the Research Council of Hong Kong.

## Author Contributions

Conceptualization, K.W.Y.Y.; Methodology, C.Y.Y.W. and K.W.Y.Y.; Data collection: C.Y.Y.W.; Data analysis, Visualization, C.Y.Y.W.; Writing, C.Y.Y.W. and K.W.Y.Y.; Supervision, Project Administration, Funding Acquisition, K.W.Y.Y.

## Materials and methods

### 1. Worm maintenance and harvesting

Hermaphrodites were maintained at room temperature (20°C) and fed on OP50 on EZ agar plates (Tris-Cl, Tris-OH, Bacto Peptone, Cholesterol, and NaCl). Worms were collected from agar plates or from the liquid culture. Embryos are harvested from hypochlorite-treated gravid-adults (1% hypochlorite, 0.5N NaOH or KOH, in M9 buffer). Lysates were incubated on ice for 10 minutes until most embryos were released. Embryos were collected by centrifuging at 700-800g for 1 minutes followed by rinsing in M9 buffer (22mM KH_2_PO_4_, 42.3mM Na_2_HPO_4_, 2mM MgSO_4_ and 85.6mM NaCl). Strains used in this study are listed in Table S1.

### 2. RNAi interference

RNAi knockdown was carried out mainly by injecting dsRNA if not stated otherwise. Other knockdown methods included feeding and soaking RNAi.

RNAi by injection: L4 larvae or young adults were injected with 1-1.5 µg/µl dsRNA of the target genes. For double knockdown, dsRNA were mixed in 1:1 ratio. For partial klp-7 RNAi, dsRNA were mixed 1:16 with MQ or with *csr-1* dsRNA as described in Gerson-Gurwitz et al. (2016). Injected worms were incubated on EZ worm plate with OP50 for 24 hours at 20-25°C before imaging or experimental procedure.

RNAi by feeding: Feeding RNAi was used when large number of embryos is needed (Figure 2C, D, F-I). L4 larvae were fed with HT115 bacteria expressing either null, hcp-3 or *csr-1* dsRNA for 24-36 hours at 20°C. The negative RNAi control was HT115 bacteria carrying empty PL4440 vector (a gift from Dr. Garry Wong lab). csr-1-dsRNA expressing bacteria was ordered from Dharmacon (CeRNAi Feeder F20D12.1). hcp-3-dsRNA vector was generated from PL4440 (WYYp142). 50 ml overnight culture was diluted in 1:50 to incubate for 3 hours at 37°C followed by 4-hours IPTG (4mM) induction. The induced bacteria were concentrated 10-fold then spiked with IPTG, ampicillin and tetracycline before seeding onto EZ agar plate. 5 μl of packed L4 larvae was added onto each worm plate with dsRNA-expressing bacteria or PL4440 containing bacteria (feeding RNAi control). About 10-15 worm plates for each treatment were used for western blotting or cDNA preparation.

RNAi by soaking: Soaking RNAi was used for ego-1(RNAi). Purified dsRNA was mixed with 10× soaking buffer (109 mM Na_2_HPO_4_, 55 mM KH_2_PO_4_, 21 mM NaCl, 47 mM NH_4_Cl, and nuclease-free H_2_O) to yield a concentration of 1 μg/μl. 20-25 L4 larvae were soaked in dsRNA solution containing 3 mM spermidine and 0.05% gelatin at 22°C for 24 hours and recovered on EZ worm plate with food at 22°C for 24 hours before analysis.

The RNAi knockdown efficiency was verified by embryonic lethality test. For those genes with no embryonic lethality upon knockdown, the knockdown efficiency was verified by RT-qPCR instead **(Figure S3B)**.

### 3. Double-stranded RNA Production

800-1,000 base pairs coding region of targeted genes was amplified from N2 *C. elegans* cDNA or genomic DNA using primers listed in Table S2. Purified PCR products were subjected to in vitro transcription (MEGAscript T3/T7 Transcription Kit; Life Technologies). Reaction products were digested with TURBO DNase at 37°C for 15 min and purified (MEGAclear Kit; Life Technologies). Eluates were incubated at 68°C for 10 minutes followed by 37°C for 30 minutes to generate double-stranded RNA (dsRNA).

### 4. Immunofluorescence

After dissection of gravid hermaphrodites, embryos were freeze-cracked in liquid nitrogen, fixed in methanol at −20°C for 30 minutes, rehydrated in PBS for 5 minutes, and blocked in AbDil (4% BSA and 0.1% Triton X-100 in PBS) at room temperature for 20 minutes. Incubation of primary antibody was done at 4°C overnight (List of antibody and condition in Table 4). Slides were washed with PBST before fluorescence-conjugated secondary antibody (1:500; Jackson ImmunoResearch Laboratories) incubation at room temperature for 1 hour. Samples was incubated with DAPI for 5 minutes and mounted (ProLong™ Diamond Antifade Mountant, Life Technologies).

### 5. Image acquisition

Confocal images were acquired from inverted confocal microscopes (LSM 710, LSM780 confocal microscope; Zeiss, VOX spinning disk confocal microscope; Pekin Elmer) with 40 or 63x oil objective and PMT detectors for DIC images. For fixed IF samples, z-stacks were captured with z-step size at 0.5 μm. Live-cell imaging was acquired at 30-second interval with a z-step interval of 1 μm. For chromatin fibers assays, images were acquired with widefield fluorescence microscope (Nikon 80i Fluorescent Microscope) with 40x 0.75 NA air objective and the imaging software (Spot Analysis). Images were processed and analyzed in the ImageJ platform.

### 6. Quantification of metaphase plate signal intensity

Metaphase fluorescence intensity was quantified as in Lee et al. (2016). Metaphase was judged as the time point right before noticeable separation of chromatin masses. The metaphase plate intensity was quantified by fitting a rectangle (A1) around the metaphase chromatin in a maximum-projected image (Figure S2A). Area of A1 were kept constant within each set of experiment. Area 2 (A2) is a slightly bigger rectangle that encloses A1. The average background intensity and the total signal intensity are calculated as below.

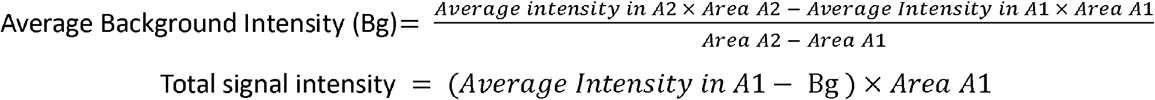

### 7. Aneuploidy assay

The aneuploidy assay was performed as described in Lee et al. (2016). The degree of cell’s aneuploidy was assessed by the chromosome number reporter strain AV221. The strain carries lacO repeats inserted in the translocated segment of Chromosome II/III and express LacI::GFP. Normal diploid embryonic cells are expected to have two GFP foci which are from the homologous chromosomes. GFP::LacI are stained by anti-GFP antibodies and foci per embryonic cells were counted from the confocal images.

### 8. Chromatin fiber preparation and analysis

The preparation of chromatin fiber was adopted from (Kyriacou and Heun, 2018) (and personal communication with Kyriacou). Chitinase-treated embryos were lysed and spread on the glass slides to a semi-dry state. Slides were incubated in lysis buffer (8M Urea and 200mM NaCl) and pulled out slowly. Rigorous lysis condition (8 M Urea) ensures removal of weakly interacting proteins but not nucleosomal proteins. Control and RNAi samples were prepared in parallel to ensure similar stretching force applied. Fibers with noticeable GFP::HCP-3, histone H4, and DAPI staining were imaged. Fibers with similar chromatin width (determined from DAPI-staining) were selected (Figure 2F-I) and images were cropped as 20-pixel-wide rectangles covering only the linear fibers for analysis. Region of interest (ROI) was defined using the ‘Analyze particles’ function in ImageJ (circularity: 0.01-1.00, exclude particles on the edges, threshold: 38 a.u.). Threshold was set based on the fiber and background signal intensities (Figure S2D). The area and the average intensities of GFP::HCP-3 and H4 staining of the foci ROI were quantified. To account for the possible variations in the degree of chromatin stretching, the GFP foci intensities were measured as a relative ratio between GFP::HCP-3 and H4. The Nearest neighbor distance of all these ROI were calculated using ‘NND’ function in ImageJ. The NND is normalized by multiplying to H4 signal for any given foci, as the more compacted chromatin is expected to give a smaller nearest neighbor distance and a higher H4 signal.

### 9. Chromatin HCP-3 turnover photobleaching assay

The photobleaching assay was performed as described in Gassmann 2012 (Figure S2C). Either untreated or *csr-1*(RNAi) OD421 (GFP::HCP-3; mCherry::H2B) hermaphrodites were dissected for live imaging using VoX spinning disk confocal microscope (Pekin Elmer). 1-cell embryos were tracked until they reached the first anaphase. At anaphase, GFP on the posterior (P-side) sister was photobleached with 440 nm laser (20%, 500 msec, 100 times). The photobleached embryo was imaged with 30-60-second interval till the metaphases of two-cell stage. The HCP-3 turnover within one cell division is calculated by the normalized GFP to mCherry intensity ratio of bleached sister chromatid (B) over that of the unbleached sister chromatid (U). For control embryos that were not photobleached, the GFP to mCherry ratio of P1 cell over AB cells were calculated.

### 10. RT-qPCR

Equal amount of untreated or RNAi-treated worms (RNAi by injection) or embryos (RNAi by feeding) were harvested for RNA extraction using standard TRIzol (Life Technologies) protocol. Reverse transcription was done using reverse-transcriptase (Applied Biosystems, 4368814). Realtime qPCR was performed in StepOne™ Real-Time PCR System using SYBR green master mix (Applied Biosystems™). Primers used for RT-qPCR are listed in Table S2.

### 11. Western blot

Embryos were lysed in 100 μl RIPA buffer using water bath sonication at 4°C for 30 minutes. Protein concentrations were determined by BCA assay (Pierce™ BCA Protein Assay Kit, ThermoFisher). 20-30 μg protein was loaded in each lane for SDS-PAGE. Proteins were transferred to PVDF membranes and blocked with 5 % milk in TBST (0.1 % Triton X-100) then probed with primary antibodies (Table S3) diluted in 5% milk in TBST at 4°C overnight. After washes with TBST, immunoblots were subjected to HRP-conjugated secondary antibodies incubation (Abcam ab97051 or ab97023) at room temperature for 1 hour followed by washing. Blot signals were developed and detected using Amersham ECL Prime western blotting detection reagent (GE Healthcare Life Sciences). Full blots of the three independent biological replicates can be found in Figure S2B.

### 12. Statistics

P-values are calculated by unpaired t-test or Fisher’s test (For frequencies, Figure1A-B). Significance would be reported if p<0.05. All error bars represent 95% confidence interval of the mean. ns: not significant; */#: p<0.05; **/##: p<0.01; ***/###: p<0.001, ****/####: p<0.0001. Graphs and plots are prepared with Microsoft excel and Prism.

## Supplemental Information

**Figure S1.**
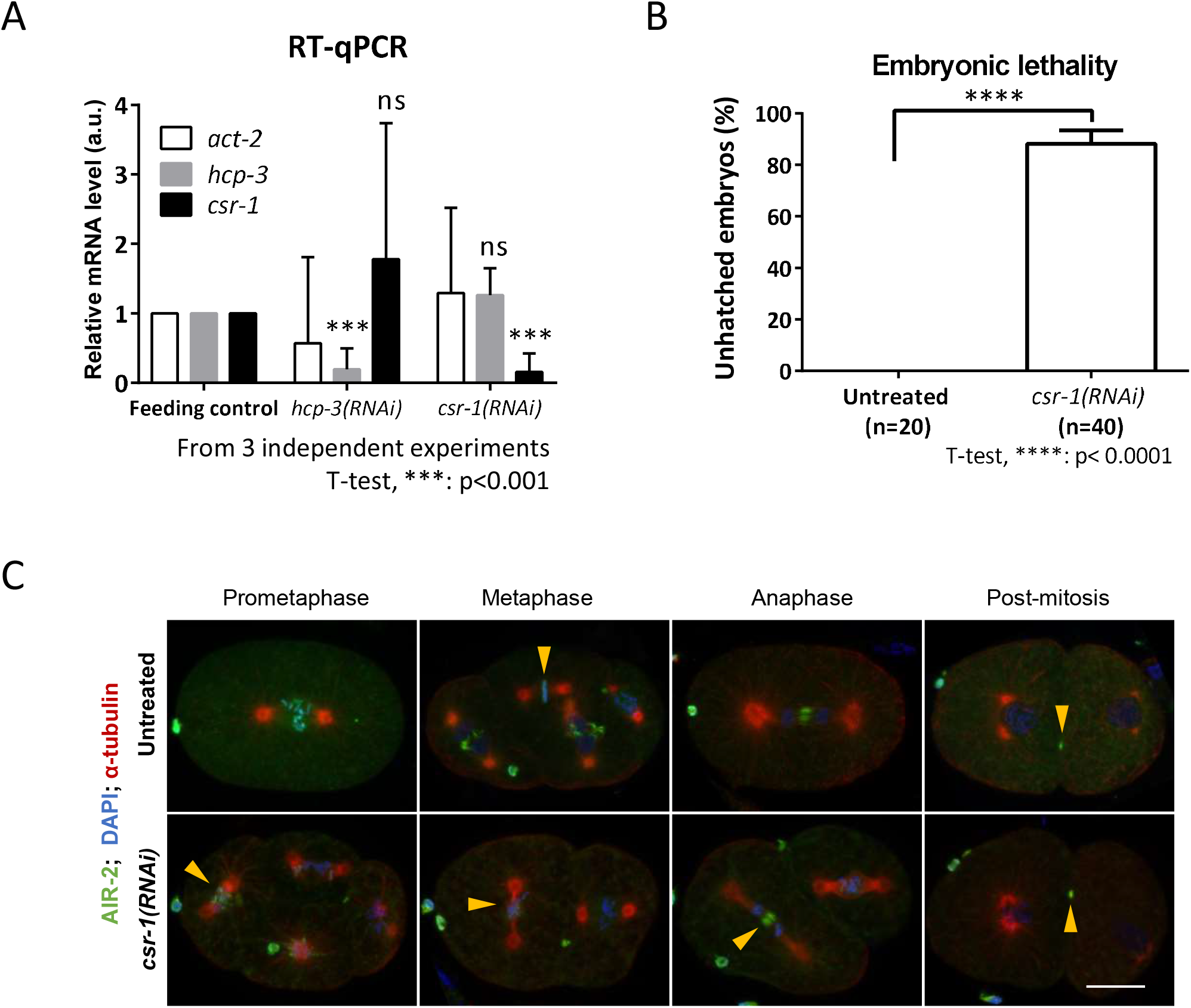
Verification of csr-1 RNA knockdown efficiency and its effect on AIR-2 localization. (A) hcp-3 and *csr-1* RNA level with hcp-3 or *csr-1*(RNAi) knockdown. act-2 was used as the loading control. Three independent biological replicates of each treatment were used. *: p<0.05, ***: p<0.001, n.s.: not significant. (B) Percentage of embryonic lethality of the untreated and RNAi knockdown worm. n equals the number of untreated/RNAi-treated animal used. (C) Immunofluorescence image showing AIR-2 localization in early embryos at different stages.

**Figure S2.**
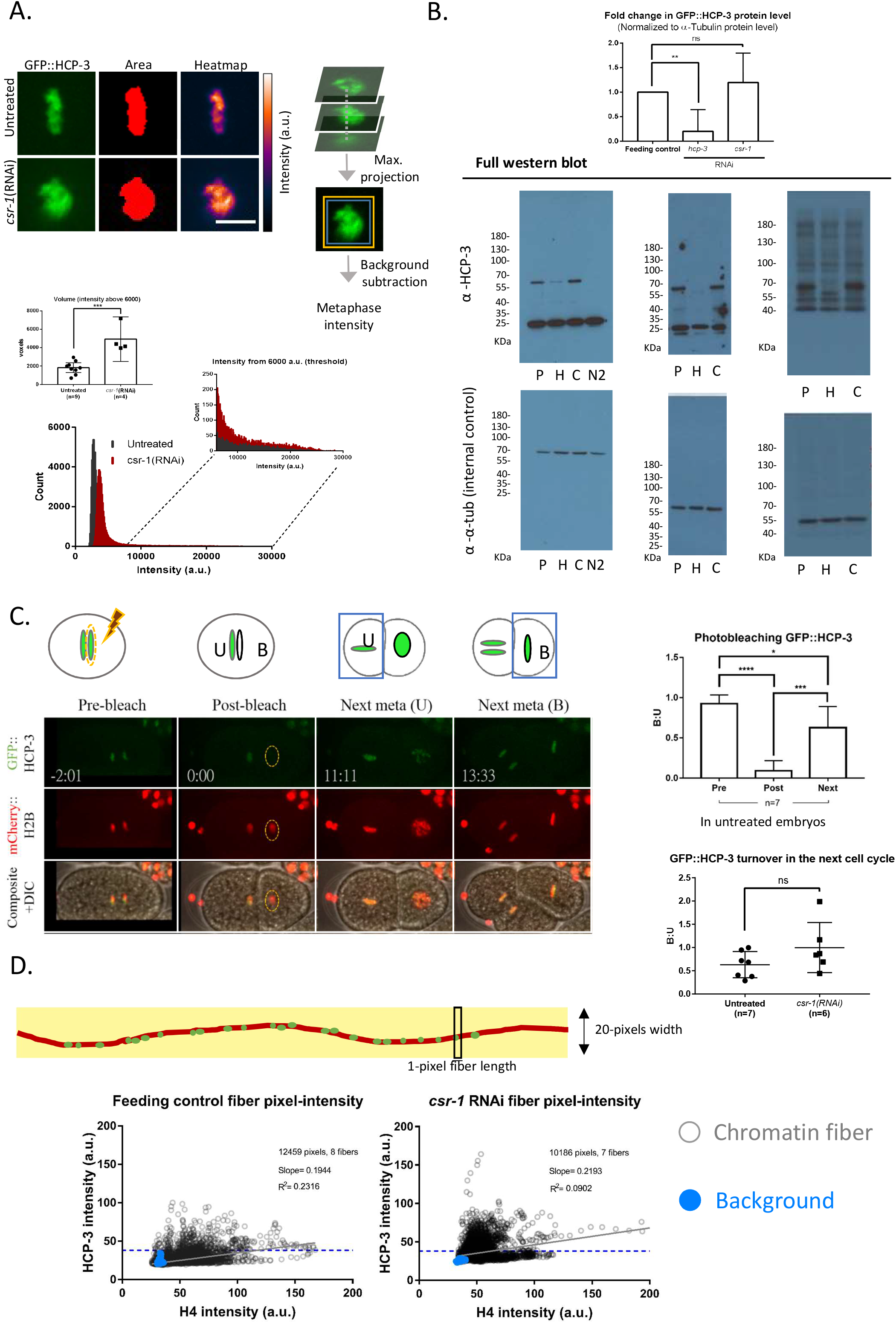
Supporting experiment details to decipher the effect of *csr-1*(RNAi) on HCP-3. (A) Metaphase chromatin in the *csr-1*(RNAi) embryos shows more high-intensity GFP::HCP-3 pixels and larger area in maximum projected image. GFP signal, area above threshold (set at 6000), and a color-coded heatmap (LUT indicated on the right) of a representative maximum projected image showing the metaphases of untreated and *csr-1*(RNAi). Illustration on the right showed how the metaphase plate total intensities were quantified in this study (See also in materials and method). Z-stack images are maximum projected. Around the chromatin, two rectangular ROIs with different size were defined. The area between the two rectangles are taken for calculating the background intensity. The mean background intensity is then subtracted from the smaller ROI. Histogram of GFP::HCP-3 intensity from the z-stacks of the representative image. An enlarged histogram showed GFP distribution above the threshold. Using the same threshold, chromatin volume of metaphase plate was quantified. n equals number of one-cell embryos quantified. (B) Full blots of the immunoblot in Figure 2E. Three biological replicates were performed. P: Feeding RNAi control (PL4440 empty vector); H: hcp-3 feeding RNAi knockdown; C: *csr-1* feeding RNAi knockdown; N2: control strain without GFP fused HCP-3. The HCP-3 level of the RNAi-treated embryos are normalized to the feeding control then to the corresponding α-tubulin level. (C) Procedure of turnover assay using photobleaching. GFP::HCP-3 on one of the anaphase sister chromatids was photobleached using high power laser with hand-drawn ROI (dashed-ellipse). The fluorescent intensities of the two daughter cells are measured in the next metaphases. The intensity ratio of the bleached over unbleached chromatin (B:U) was measured to quantify protein turnover on chromatin. The reappeared GFP signal on the bleached chromatin represents newly assembled GFP::HCP-3, which is compared to the total HCP-3 on chromatin as observed in the unbleached chromatin. The quantification of the ROI GFP signal before and after photobleaching as a fold change to the unbleached sister. The B:U for untreated control and *csr-1*(RNAi). n equals the number of photobleached embryos analyzed. (D) Scatter plots of the GFP::HCP-3 and H4 pixel intensity of the pixels on the chromatin fibers prepared from the control (8 fibers) and the *csr-1*(RNAi) embryos (7 fibers). Segment lines (20-pixel-wide) are drawn along the image of fibers. The mean-intensity of these 20 pixels was determined. The linear regression line and the R-square values were indicated. Blue data points are the pixels of a line drawn at non-fiber background in the fluorescent images. 1533 pixels of fiber -length have been sampled for feeding RNAi control and 1581 pixels for csr-1(RNAi) from the background. A threshold for GFP::HCP-3 intensity was selected (38 a.u., blue dashed lines) to exclude fiber pixels with HCP-3 intensity close to the background level for both the feeding control and the csr-1(RNAi) embryos.

**Figure S3.**
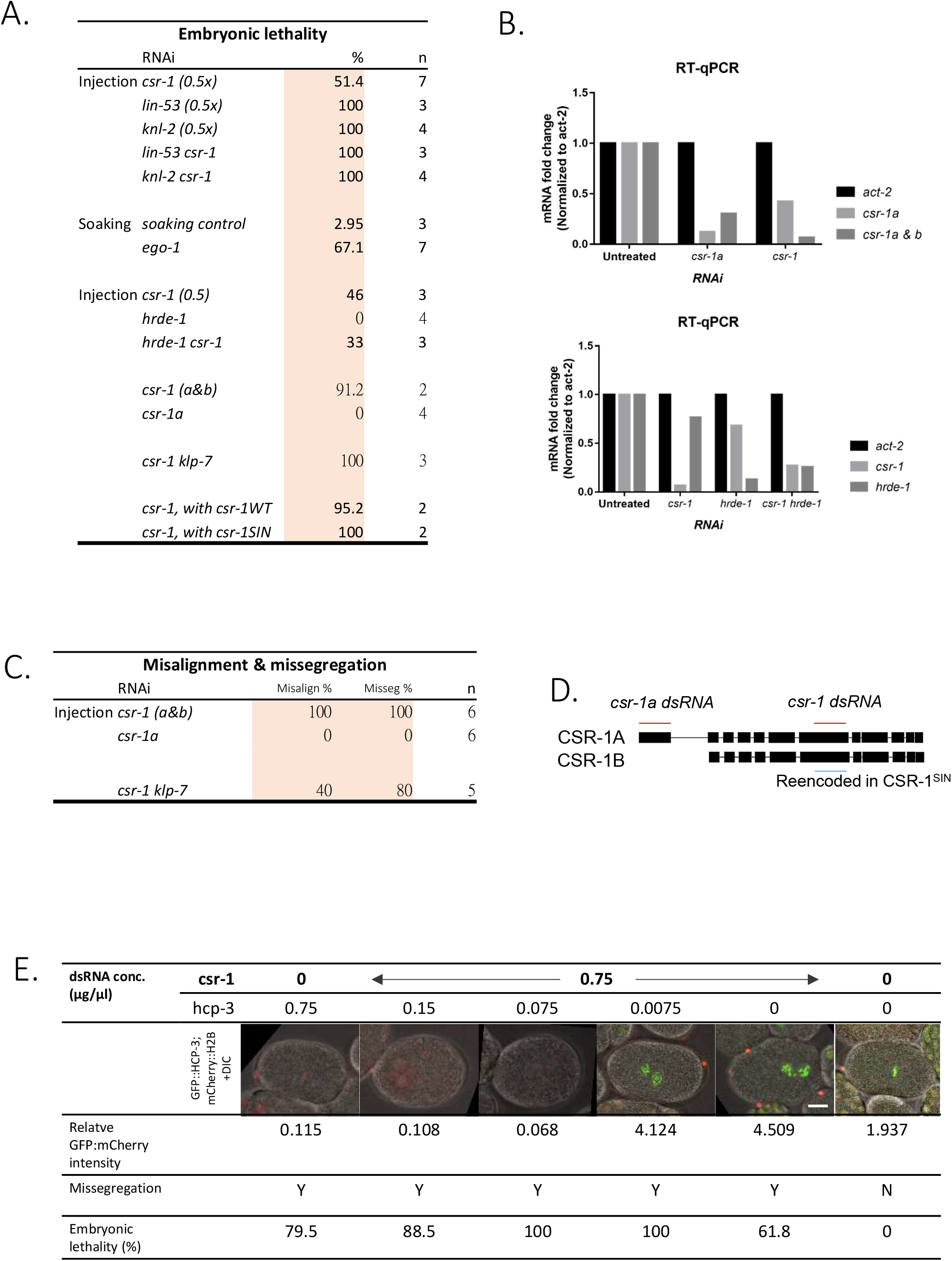
Verification of RNAi knockdown efficiency, transgene expression, segregation fate and embryonic lethality. (A) Embryonic lethality of different RNAi treatments in Figure 3. Hermaphrodites were isolated for laying embryos from 24 hours post-RNAi treatment. Percentage of unhatched embryos for each hermaphrodite was quantified. Average percentage was shown and n equals number of hermaphrodites used. (B) Quantification of RNA levels of *csr-1* isoforms, hrde-1 and act-2 by RT-qPCR in hrde-1(RNAi) and *csr-1*a(RNAi) worms or double RNAi-treated worms. 2 technical qPCR replicates were performed, and the mean of the 2 replicates were plotted. (C) Frequency of misalignment and missegregation in *csr-1* klp-7(RNAi) knockdown embryos. (D) Gene annotation of two *csr-1* isoforms. Positions for dsRNA design to knockdown *csr-1* and specifically knockdown *csr-1*a are in red. The region reencoded in the CSR-1 mutant (WT/SIN) is highlighted in blue. Note that the reencoded region overlaps with the *csr-1* dsRNA, so the mutants used in Figure 3F are RNAi-resistant. (E) HCP-3 partial knockdown in csr-1 (RNAi), to observe if reducing HCP-3 level can rescue chromosome missegregation phenotype. The GFP::HCP-3 and mCherry::H2B signals are shown and embryonic lethality frequency is shown.

**Table S1.**
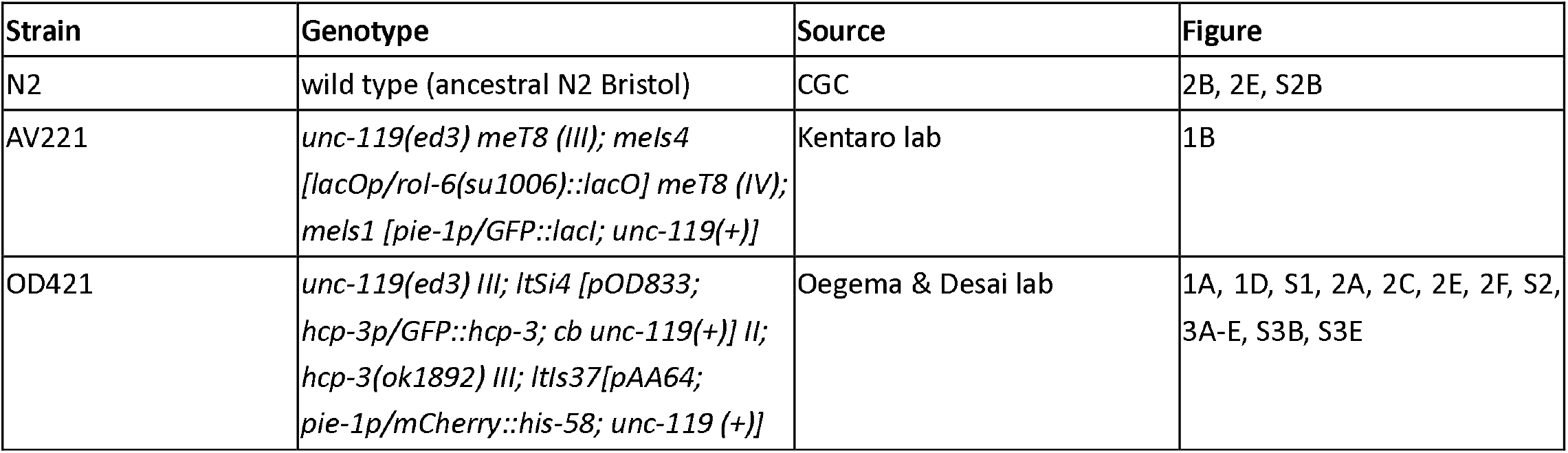
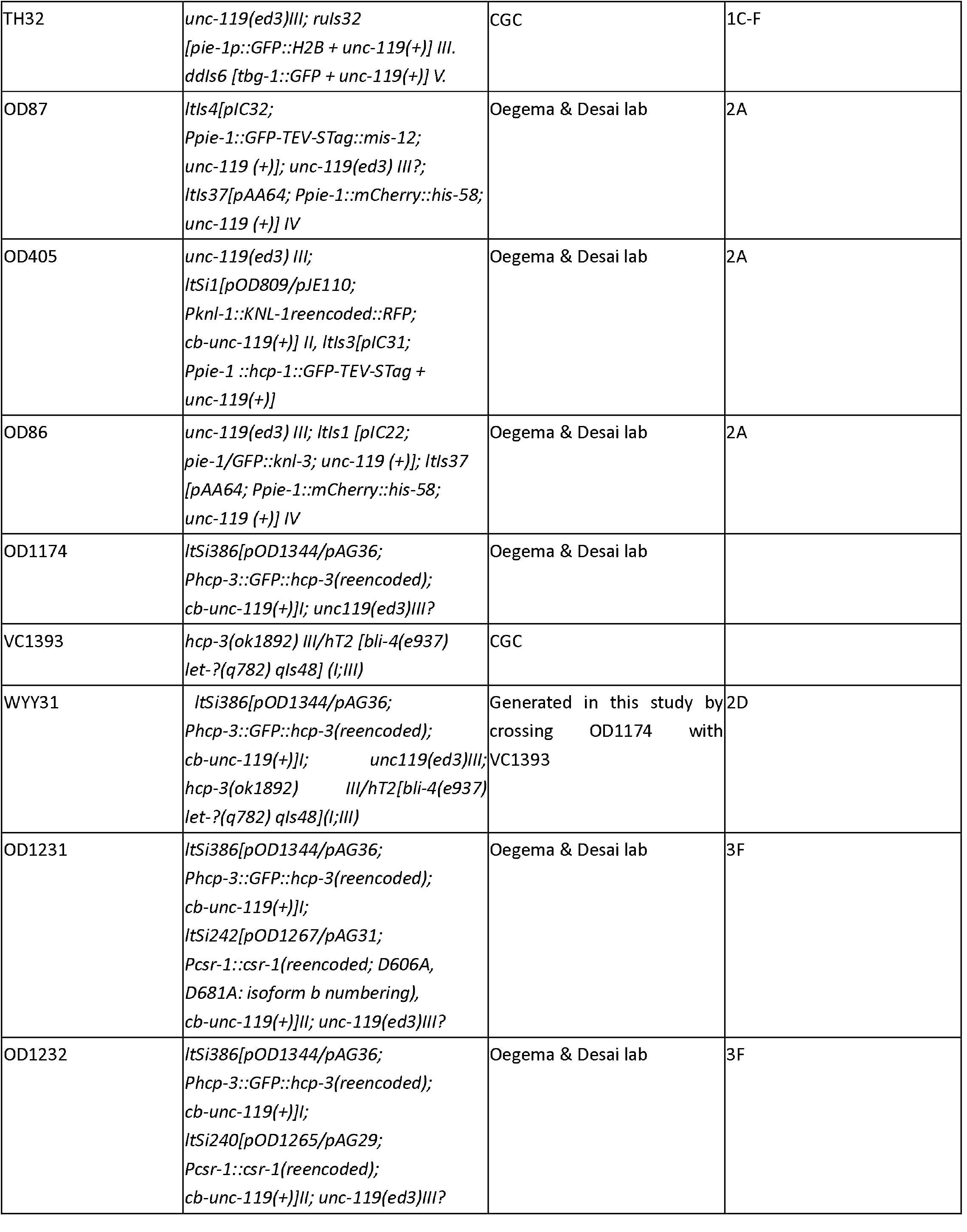
Strain used in this study.

**Table S2.**
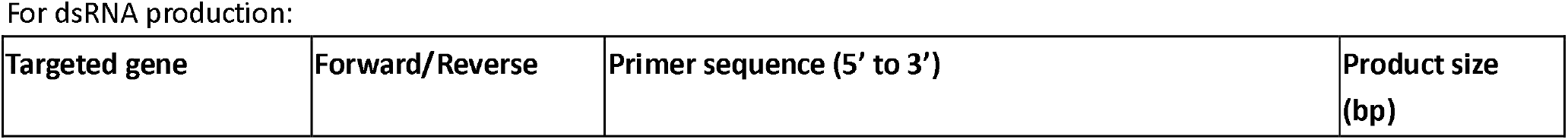
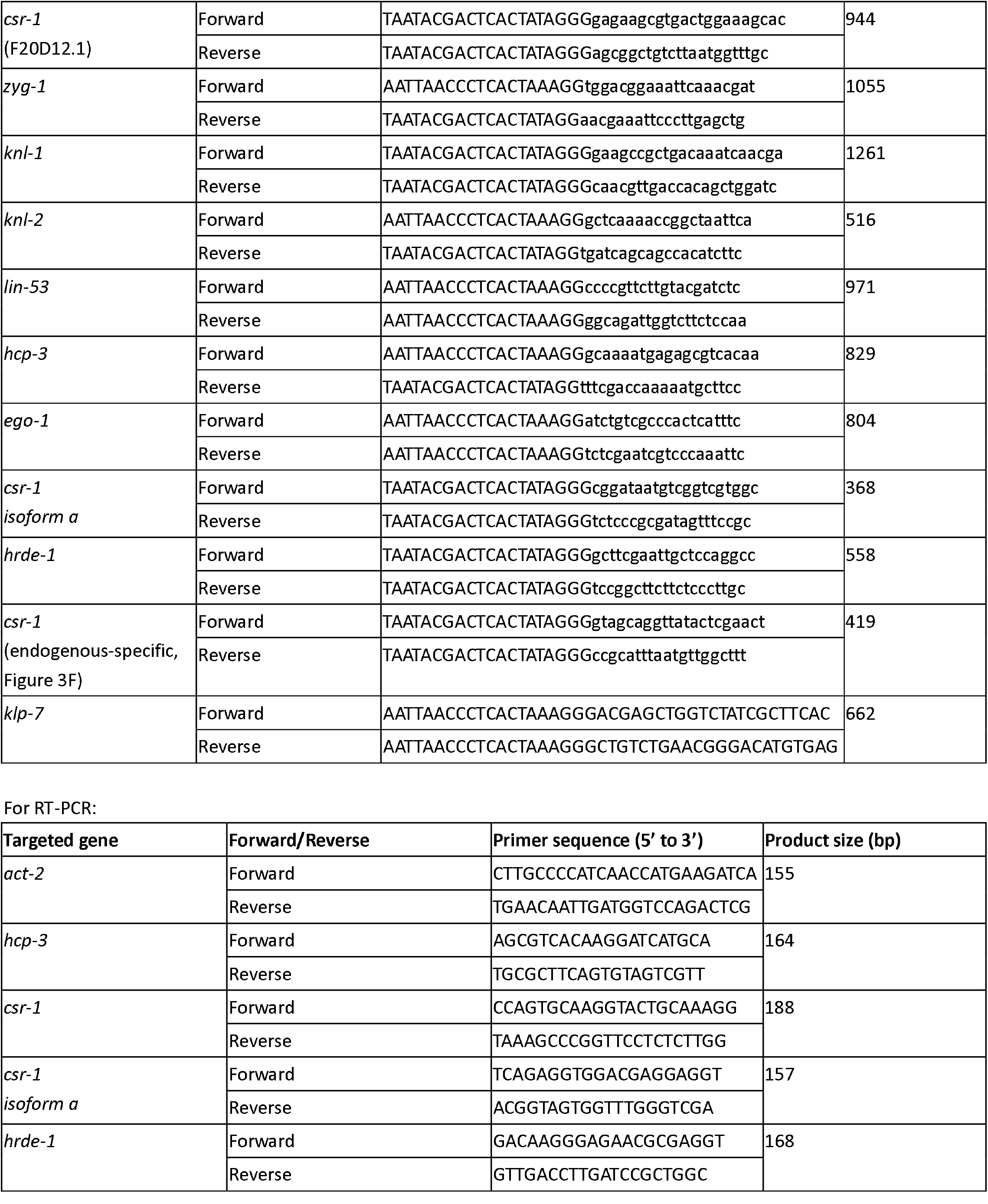
Oligo used in this study.

**Table S3.** Antibody list.

| Antibody (against) | Host | Dilution | Source | Used in Figure |
| --- | --- | --- | --- | --- |
| GFP | Rabbit | 1:1000 | Novus (NB600-308) | 1A, 1B, 2E |
| α-Tubulin | Mouse | 1:3000 | Abcam (ab7291) | 1A, S1C, 2E |
| LacI | Mouse | 1:500 | Millipore (05-503) | 1B |
| AIR-2 | Rabbit | 1:1000 | Gift from OD lab | S1C |
| HCP-3 | Rabbit | 1:1000 | Novus (29540002) | 2B |
| KNL-2 | Rabbit | 1:1000 | Gift from OD lab | 2B |
| NDC-80 | Rabbit | 1:1000 | Gift from OD lab | 2B |
| HCP-4 | Rabbit | 1:1000 | Gift from OD lab | 2B |
| MIS-12 | Rabbit | 1:1000 | Novus (35550002) | 2B |
| Histone H4 | Rabbit | 1:500 | Abcam (ab10158) | 2F |
| GFP | Mouse | 1:500 | Abcam (ab1218) | 2F |

## Graphical abstract

CSR-1 is a repressor for HCP-3 chromatin localization

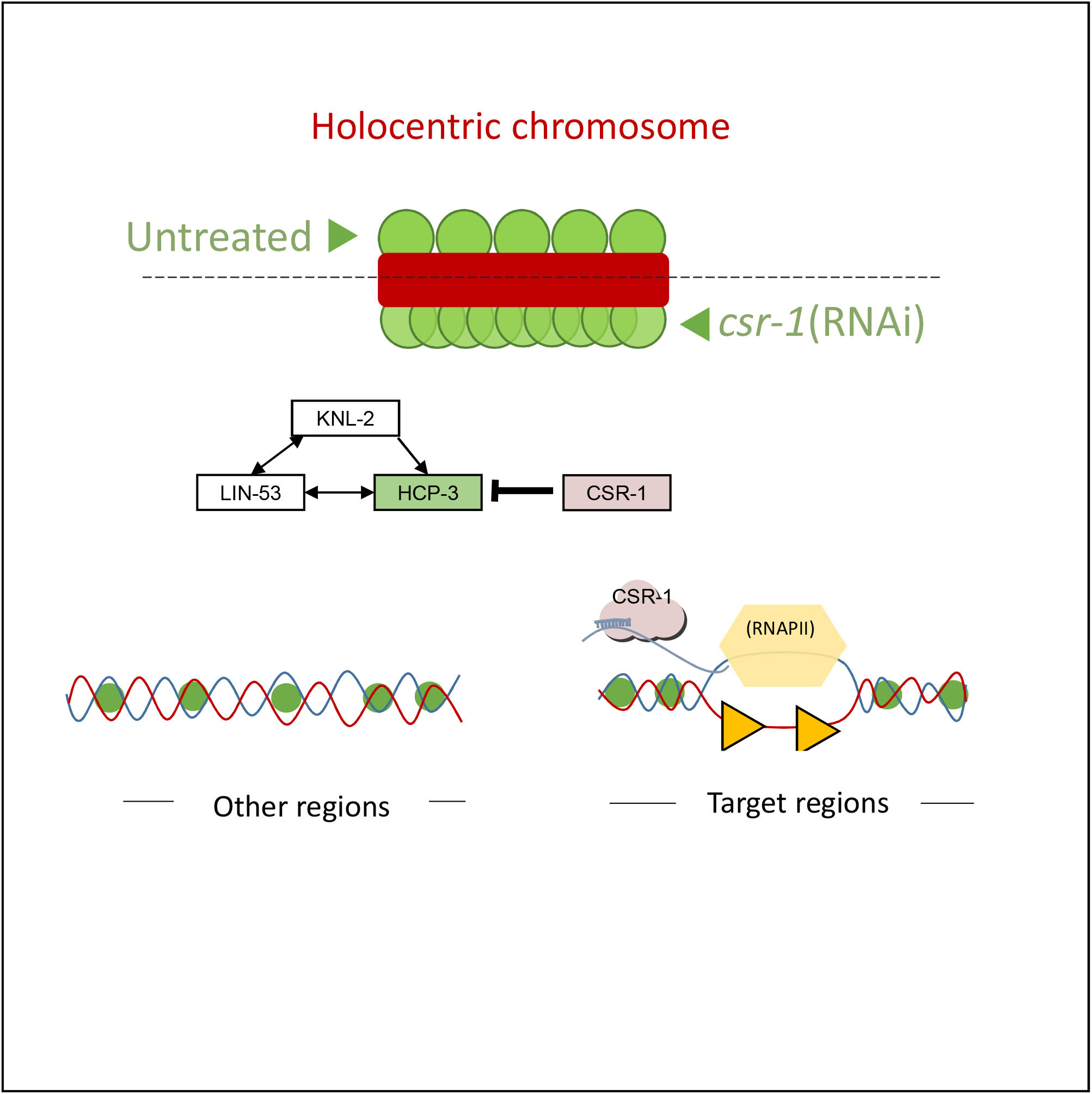

## References

Amato, A., Schillaci, T., Lentini, L. & Di Leonardo, A. 2009. CENPA overexpression promotes genome instability in pRb-depleted human cells. Molecular cancer, 8, 119.

Amor, D. J. & Choo, K. A. 2002. Neocentromeres: role in human disease, evolution, and centromere study. The American Journal of Human Genetics, 71, 695–714.

Antonio, C., Ferby, I., Wilhelm, H., Jones, M., Karsenti, E., Nebreda, A. R. & Vernos, I. 2000. Xkid, a chromokinesin required for chromosome alignment on the metaphase plate. Cell, 102, 425–435.

Aoki, K., Moriguchi, H., Yoshioka, T., Okawa, K. & Tabara, H. 2007. In vitro analyses of the production and activity of secondary small interfering RNAs in C. elegans. The EMBO journal, 26, 5007–5019.

Avgousti, D. C., Palani, S., Sherman, Y. & Grishok, A. 2012. CSR-1 RNAi pathway positively regulates histone expression in C. elegans. The EMBO journal, 31, 3821–3832.

Bajer, A. S. & Molø-Bajer, J. 1972. Spindle dynamics and chromosome movements. International review of cytology.

Buchwitz, B. J., Ahmad, K., Moore, L. L., Roth, M. B. & Henikoff, S. 1999. A histone-H3-like protein in C. elegans. Nature, 401, 547–8.

Caron, C., Kemphues, K. & White, J. The zyg-1 Gene Encodes a Serine/Threonine Kinase Required for Bipolar Spindle Formation. International C. elegans Meeting, 1999.

Castillo, A. G., Mellone, B. G., Partridge, J. F., Richardson, W., Hamilton, G. L., Allshire, R. C. & Pidoux, A. L. 2007. Plasticity of fission yeast CENP-A chromatin driven by relative levels of histone H3 and H4. PLoS genetics, 3, e121.

Charlesworth Amanda G., Seroussi, U., Lehrbach Nicolas J., Renaud Mathias S., Sundby Adam E., Molnar Ruxandra I., Lao Robert X., Willis Alexandra R., Woock Jenna R., Aber Matthew J., Diao Annette J., Reinke Aaron W., Ruvkun, G. & Claycomb Julie M. 2021. Two isoforms of the essential C. elegans Argonaute CSR-1 differentially regulate sperm and oocyte fertility. Nucleic Acids Research, 49, 8836–8865.

Cimini, D., Howell, B., Maddox, P., Khodjakov, A., Degrassi, F. & Salmon, E. 2001. Merotelic kinetochore orientation is a major mechanism of aneuploidy in mitotic mammalian tissue cells. The Journal of cell biology, 153, 517–528.

Claycomb, J. M., Batista, P. J., Pang, K. M., Gu, W., Vasale, J. J., Van Wolfswinkel, J. C., Chaves, D. A., Shirayama, M., Mitani, S., Ketting, R. F., Conte, D. & Mello, C. C. 2009. The Argonaute CSR-1 and its 22G-RNA cofactors are required for holocentric chromosome segregation. Cell, 139, 123–34.

De Rop, V., Padeganeh, A. & Maddox, P. S. 2012. CENP-A: the key player behind centromere identity, propagation, and kinetochore assembly. Chromosoma, 121, 527–38.

Dumont, J., Oegema, K. & Desai, A. 2010. A kinetochore-independent mechanism drives anaphase chromosome separation during acentrosomal meiosis. Nature cell biology, 12, 894–901.

Essex, A., Dammermann, A., Lewellyn, L., Oegema, K. & Desai, A. 2009. Systematic Analysis in Caenorhabditis elegans Reveals that the Spindle Checkpoint Is Composed of Two Largely Independent Branches. Molecular Biology of the Cell, 20, 1252–1267.

Eulalio, A., Huntzinger, E. & Izaurralde, E. 2008. GW182 interaction with Argonaute is essential for miRNA-mediated translational repression and mRNA decay. Nature structural & molecular biology, 15, 346.

Friend, K., Campbell, Z. T., Cooke, A., Kroll-Conner, P., Wickens, M. P. & Kimble, J. 2012. A conserved PUF–Ago–eEF1A complex attenuates translation elongation. Nature Structural and Molecular Biology, 19, 176.

Gassmann, R., Rechtsteiner, A., Yuen, K. W., Muroyama, A., Egelhofer, T., Gaydos, L., Barron, F., Maddox, P., Essex, A., Monen, J., Ercan, S., Lieb, J. D., Oegema, K., Strome, S. & Desai, A. 2012. An inverse relationship to germline transcription defines centromeric chromatin in C. elegans. Nature, 484, 534–7.

Gerson-Gurwitz, A., Wang, S., Sathe, S., Green, R., Yeo, G. W., Oegema, K. & Desai, A. 2016. A small RNA-catalytic Argonaute pathway tunes germline transcript levels to ensure embryonic divisions. Cell, 165, 396–409.

Gerstein, M. B., Lu, Z. J., Van Nostrand, E. L., Cheng, C., Arshinoff, B. I., Liu, T., Yip, K. Y., Robilotto, R., Rechtsteiner, A., Ikegami, K., Alves, P., Chateigner, A., Perry, M., Morris, M., Auerbach, R. K., Feng, X., Leng, J., Vielle, A., Niu, W., Rhrissorrakrai, K., Agarwal, A., Alexander, R. P., Barber, G., Brdlik, C. M., Brennan, J., Brouillet, J. J., Carr, A., Cheung, M. S., Clawson, H., Contrino, S., Dannenberg, L. O., Dernburg, A. F., Desai, A., Dick, L., Dose, A. C., Du, J., Egelhofer, T., Ercan, S., Euskirchen, G., Ewing, B., Feingold, E. A., Gassmann, R., Good, P. J., Green, P., Gullier, F., Gutwein, M., Guyer, M. S., Habegger, L., Han, T., Henikoff, J. G., Henz, S. R., Hinrichs, A., Holster, H., Hyman, T., Iniguez, A. L., Janette, J., Jensen, M., Kato, M., Kent, W. J., Kephart, E., Khivansara, V., Khurana, E., Kim, J. K., Kolasinska-Zwierz, P., Lai, E. C., Latorre, I., Leahey, A., Lewis, S., Lloyd, P., Lochovsky, L., Lowdon, R. F., Lubling, Y., Lyne, R., Maccoss, M., Mackowiak, S. D., Mangone, M., Mckay, S., Mecenas, D., Merrihew, G., Miller, D. M., 3rd, Muroyama, A., Murray, J. I., Ooi, S. L., Pham, H., Phippen, T., Preston, E. A., Rajewsky, N., Ratsch, G., Rosenbaum, H., Rozowsky, J., Rutherford, K., Ruzanov, P., Sarov, M., Sasidharan, R., Sboner, A., Scheid, P., Segal, E., Shin, H., Shou, C., Slack, F. J., et al. 2010. Integrative analysis of the Caenorhabditis elegans genome by the modENCODE project. Science, 330, 1775–87.

Gregan, J., Polakova, S., Zhang, L., ToliĆ-N Trends in cell biology, 21, 374–381.

Grill, S. W., Goènczy, P., Stelzer, E. H. & Hyman, A. A. 2001. Polarity controls forces governing asymmetric spindle positioning in the Caenorhabditis elegans embryo. Nature, 409, 630–633.

Gu, W., Shirayama, M., Conte, D., Vasale, J., Batista, P. J., Claycomb, J. M., Moresco, J. J., Youngman, E. M., Keys, J. & Stoltz, M. J. 2009. Distinct argonaute-mediated 22G-RNA pathways direct genome surveillance in the C. elegans germline. Molecular cell, 36, 231–244.

Guenatri, M., Bailly, D., Maison, C. & Almouzni, G. 2004. Mouse centric and pericentric satellite repeats form distinct functional heterochromatin. Journal of Cell Biology, 166, 493–505.

Guo, Y., Kim, C. & Mao, Y. 2013. New insights into the mechanism for chromosome alignment in metaphase. International review of cell and molecular biology. Elsevier.

Heun, P., Erhardt, S., Blower, M. D., Weiss, S., Skora, A. D. & Karpen, G. H. 2006. Mislocalization of the Drosophila Centromere-Specific Histone CID Promotes Formation of Functional Ectopic Kinetochores. Dev Cell, 10, 303–15.

Huang, C., Wang, X., Liu, X., Cao, S. & Shan, G. 2015. RNAi pathway participates in chromosome segregation in mammalian cells. Cell Discovery, 1, 15029.

J. Richard Mcintosh, Ekaterina L. Grishchuk & West, R. R. 2002. Chromosome-Microtubule Interactions During Mitosis. Annual Review of Cell and Developmental Biology, 18, 193–219.

Kaitna, S., Pasierbek, P., Jantsch, M., Loidl, J. & Glotzer, M. 2002. The aurora B kinase AIR-2 regulates kinetochores during mitosis and is required for separation of homologous chromosomes during meiosis. Current biology, 12, 798–812.

Kyriacou, E. & Heun, P. 2018. High-resolution mapping of centromeric protein association using APEX-chromatin fibers. Epigenetics & chromatin, 11, 1–17.

Ladouceur, A.-M., Ranjan, R., Smith, L., Fadero, T., Heppert, J., Goldstein, B., Maddox, A. S. & Maddox, P. S. 2017. CENP-A and topoisomerase-II antagonistically affect chromosome length. J Cell Biol, 216, 2645–2655.

Lee, B. C. H., Lin, Z. & Yuen, K. W. Y. 2016. RbAp46/48 LIN-53 Is Required for Holocentromere Assembly in Caenorhabditis elegans. Cell reports, 14, 1819–1828.

Lin, Z., Xie, Y., Nong, W., Ren, X., Li, R., Zhao, Z., Hui Jerome Ho L. & Yuen Karen Wing Y. 2021. Formation of artificial chromosomes in Caenorhabditis elegans and analyses of their segregation in mitosis, DNA sequence composition and holocentromere organization. Nucleic Acids Research, 49, 9174–9193.

Liu, J., Carmell, M. A., Rivas, F. V., Marsden, C. G., Thomson, J. M., Song, J.-J., Hammond, S. M., Joshua-Tor, L. & Hannon, G. J. 2004. Argonaute2 is the catalytic engine of mammalian RNAi. Science, 305, 1437–1441.

Liu, J., Rivas, F. V., Wohlschlegel, J., Yates, J. R., Parker, R. & Hannon, G. J. 2005. A role for the P-body component GW182 in microRNA function. Nature cell biology, 7, 1261–1266.

Maddox, P. S., Hyndman, F., Monen, J., Oegema, K. & Desai, A. 2007. Functional genomics identifies a Myb domain–containing protein family required for assembly of CENP-A chromatin. The Journal of Cell Biology, 176, 757–763.

Mcewen, B. F., Hsieh, C. E., Mattheyses, A. L. & Rieder, C. L. 1998. A new look at kinetochore structure in vertebrate somatic cells using high-pressure freezing and freeze substitution. Chromosoma, 107, 366–75.

Oegema, K., Desai, A., Rybina, S., Kirkham, M. & Hyman, A. A. 2001. Functional analysis of kinetochore assembly in Caenorhabditis elegans. The Journal of cell biology, 153, 1209–1226.

Powers, J., Rose, D. J., Saunders, A., Dunkelbarger, S., Strome, S. & Saxton, W. M. 2004. Loss of KLP-19 polar ejection force causes misorientation and missegregation of holocentric chromosomes. J Cell Biol, 166, 991–1001.

Redemann, S., Baumgart, J., Lindow, N., Shelley, M., Nazockdast, E., Kratz, A., Prohaska, S., Brugu¢s, J., FNat Commun, 8, 15288.

Schwarz, D. S., Tomari, Y. & Zamore, P. D. 2004. The RNA-induced silencing complex is a Mg2+-dependent endonuclease. Current Biology, 14, 787–791.

Seth, M., Shirayama, M., Gu, W., Ishidate, T., Conte, D. & Mello, C. C. 2013. The C. elegans CSR-1 argonaute pathway counteracts epigenetic silencing to promote germline gene expression. Developmental cell, 27, 656–663.

Singh, M., Cornes, E., Li, B., Quarato, P., Bourdon, L., Dingli, F., Loew, D., Proccacia, S. & Cecere, G. 2021. Translation and codon usage regulate Argonaute slicer activity to trigger small RNA biogenesis. Nature Communications, 12, 3492.

Srayko, M., Kaya, A., Stamford, J. & Hyman, A. A. 2005. Identification and characterization of factors required for microtubule growth and nucleation in the early C. elegans embryo. Dev Cell, 9, 223–36.

Steiner, F. A. & Henikoff, S. 2014. Holocentromeres are dispersed point centromeres localized at transcription factor hotspots. Elife, 3, e02025.

Sullivan, L. L., Chew, K. & Sullivan, B. A. 2017. α satellite DNA variation and function of the human centromere. Nucleus, 8, 331–339.

Tanaka, T. U. 2002. Bi-orienting chromosomes on the mitotic spindle. Current opinion in cell biology, 14, 365–371.

Tomonaga, T., Matsushita, K., Yamaguchi, S., Oohashi, T., Shimada, H., Ochiai, T., Yoda, K. & Nomura, F. 2003. Overexpression and mistargeting of centromere protein-A in human primary colorectal cancer. Cancer research, 63, 3511–3516.

Tu, S., Wu, M. Z., Wang, J., Cutter, A. D., Weng, Z. & Claycomb, J. M. 2015. Comparative functional characterization of the CSR-1 22G-RNA pathway in Caenorhabditis nematodes. Nucleic Acids Res, 43, 208–24.

Van Wolfswinkel, J. C., Claycomb, J. M., Batista, P. J., Mello, C. C., Berezikov, E. & Ketting, R. F. 2009. CDE-1 affects chromosome segregation through uridylation of CSR-1-bound siRNAs. Cell, 139, 135–148.

Wedeles, C. J., Wu, M. Z. & Claycomb, J. M. 2013a. A multitasking Argonaute: exploring the many facets of C. elegans CSR-1. Chromosome Research, 21, 573–586.

Wedeles, C. J., Wu, M. Z. & Claycomb, J. M. 2013b. Protection of germline gene expression by the C. elegans Argonaute CSR-1. Developmental cell, 27, 664–671.

Xu, F., Feng, X., Chen, X., Weng, C., Yan, Q., Xu, T., Hong, M. & Guang, S. 2018. A cytoplasmic Argonaute protein promotes the inheritance of RNAi. Cell reports, 23, 2482–2494.

Yigit, E., Batista, P. J., Bei, Y., Pang, K. M., Chen, C.-C. G., Tolia, N. H., Joshua-Tor, L., Mitani, S., Simard, M. J. & Mello, C. C. 2006. Analysis of the C. elegans Argonaute family reveals that distinct Argonautes act sequentially during RNAi. Cell, 127, 747–757.

